# How predictability and individual alpha frequency shape memory: insights from an event-related potential investigation

**DOI:** 10.1101/2023.10.25.564081

**Authors:** Sophie Jano, Alex Chatburn, Zachariah Cross, Matthias Schlesewsky, Ina Bornkessel-Schlesewsky

## Abstract

Prediction and memory are strongly intertwined, with predictions relying on memory retrieval, whilst also influencing memory encoding. However, it is unclear how predictability influences explicit memory performance, and how individual neural factors may modulate this relationship. The current study sought to investigate the effect of predictability on memory processing with an analysis of the N400 event-related potential in a context extending beyond language. Participants (*N* = 48, females = 33) completed a study-test paradigm where they first viewed predictable and unpredictable four-item ‘ABCD’ sequences of outdoor scene images, whilst their brain activity was recorded using electroencephalography (EEG). Subsequently, their memory for the images was tested, and N400 patterns during learning were compared with memory outcomes. Behavioural results revealed better memory for images in predictable sequences in contrast to unpredictable sequences. Memory was also strongest for predictable images in the ‘B’ position, suggesting that when processing longer sequences, the brain may prioritise the data deemed most informative. Strikingly, greater N400 amplitudes during learning were associated with enhanced memory at test for individuals with low versus high individual alpha frequencies (IAFs). In light of the relationship between the N400 and stimulus predictability, this finding may imply that predictive processing differs between individuals to influence the extent of memory encoding. Finally, exploratory analyses provided evidence for a later positivity that was predictive of subsequent memory performance. Ultimately, the results highlight the complex and interconnected relationship between predictive processing and memory, whilst shedding light on the accumulation of predictions across longer sequences.

## 1. Introduction

Modern conceptualisations of memory highlight its importance for predicting the future as opposed to purely remembering the past (Bar, 2007; Exton-McGuinness et al., 2015). According to predictive coding, the brain generates predictions about future inputs from prior experience, which are refined in response to prediction errors (mismatches between the prediction and the incoming sensory input; Friston, 2010; Friston & Kiebel, 2009; Hohwy, 2013; Rao & Ballard, 1999). This refinement optimises predictions towards a maximisation of explanatory power, such that the causes of sensory inputs are more accurately inferred and error (or free energy) in the system is reduced (Friston, 2009, 2010). However, the consequences of prediction for learning and memory are not entirely evident, with the relationship between memory and predictive processing stemming from a reliance of predictions on past experiences (involving the retrieval of established memory representations; Chow et al., 2016; Rauss & Born, 2017), and from the effect of prediction on information uptake (concerning memory encoding; Kim et al., 2020; Sherman & Turk-Browne, 2020). Therefore, not only do predictions appear to rely on memory, but they may have implications for informational accuracy and biases. The present experiment sought to shed light on the effect of predictability on memory performance via an investigation into the neural underpinnings of predictive processing across sequences.

### 1.1. Predictions rely on memories of statistical relationships

The notion that predictions rely on memory representations is grounded in the brain’s ability to extract environmental regularities – a process known as statistical learning (Turk-Browne in Dodd & Flowers, 2012; Siegelman & Frost, 2015). Regularities are abundant in the natural world, such that similar patterns are encountered across time and space (Turk-Browne in Dodd & Flowers, 2012, p. 118). There is strong evidence that the brain can learn and encode patterns of this nature into memory (i.e., that it can engage in statistical learning); for example, classic visual statistical learning tasks show successful learning of triplets of shapes (Fiser & Aslin, 2002) and syllables (Saffran et al., 1996), presented in a continuous sequence (for similar research in mice, see Gavornik & Bear, 2014). This learning also appears relatively implicit, with participants judging patterns as more familiar than random shapes despite being unaware of an underlying relationship and having low confidence in their decisions (Turk-Browne in Dodd & Flowers, 2012, p. 123). This proficiency for pattern learning has been shown to facilitate predictive processing, with research demonstrating enhanced prediction of upcoming visual orientations following structured versus random sequences (Baker et al., 2014). Additionally, studies employing Serial Reaction Time tasks have revealed faster reaction times for spatial locations that were preceded by predictable versus unpredictable sequences (Nissen & Bullemer, 1987, in Schwarb & Schumacher, 2012). As such, a statistical learning mechanism within memory has been proposed, where statistical relationships are encoded as generalized memory traces to enable the anticipation of future states (Bar, 2007; Turk-Browne in Dodd & Flowers, 2012; Sherman & Turk-Browne, 2020). Statistical learning may thus act as a precursor to the formation of a predictive model (Rauss & Born, 2017), such that regularities are learnt and stored as associative traces in memory, which are subsequently retrieved for future use (Bar, 2007). This highlights the reliance of prediction on memory, with both prediction and episodic recall proposed to share hippocampal machinery (Barron et al., 2020). This common hippocampal dependence, coupled with the brain’s ability to encode environmental regularities into memory, places statistical learning and the retrieval of statistical regularities from memory, as likely prerequisites for predictive processing.

### 1.2. Prediction influences memory encoding

The dependence of both memory and prediction on the hippocampus may additionally explain the role of prediction in regulating memory encoding. Prior research has demonstrated poorer memory for stimuli that are predictive of upcoming items, suggesting that the brain may be unable to predict information and encode new memories simultaneously (Sherman & Turk-Browne, 2020). According to Sherman and Turk-Browne (2020), prediction may expend hippocampal memory retrieval resources that cannot be concurrently used to encode new traces. In the face of a prediction violation, the hippocampus is proposed to switch to an encoding state to incorporate the sensory information (Mack et al., 2018; Sinclair et al., 2021). This suggests that prediction errors may drive the formation of new memories; a proposal that is supported by behavioural research demonstrating enhanced memory for inconsistent (high prediction error-eliciting) items (Greve et al., 2017), reward prediction errors (Jang et al., 2019; Rouhani et al., 2018), surprising images (Michelon et al., 2003), bizarre sentences (Worthen & Roark, 2002), novel words (Habib et al., 2003), and unpredictable, implausible nouns (however, see Haeuser & Kray, 2022 for a discussion of this effect as being driven by the implausibility of the word, as opposed to the predictability itself). To further support the suggestion that the brain prioritises unpredictable information, Furutachi and colleagues (2023) observed amplified sensory representations in the visual cortex of mice for prediction error-eliciting stimuli. According to predictive coding perspectives, prediction errors represent sensory data (Feldman & Friston, 2010). Consequently, amplified sensory representations (observed in Furutachi et al., 2023) may imply stronger prediction errors corresponding to greater model revision and information uptake. As such, the prediction error signals resulting from expectation violations may disrupt predictive processing, such that the brain switches to an encoding state to prioritise updating of the incoming information into the predictive model. Naturally, such a mechanism would likely have important consequences for everyday functioning, as novel information could be prioritised during memory encoding.

Although prediction error-eliciting information may be preferentially encoded into memory, other research demonstrates better memory for stimuli eliciting low prediction error (i.e., predictable information; Turan et al., 2023), or for both highly expected and unexpected stimuli, in contrast to moderately expected information (Quent et al., 2022). This has prompted the suggestion that the relationship between predictability and memory follows a U-shaped, nonlinear pattern, such that schema-congruent and schema-incongruent information is prioritised in memory (with schemas proposed to form the basis of predictions; Quent et al., 2022; Van Kesteren et al., 2012). On a similar note, Bein and colleagues (2023) propose that consistency with expectations promotes the integration of that information into existing memory representations. However, prediction errors can promote both integration and memory separation, depending on a number of factors (e.g., the type of prediction error and the strength of the memory reactivation associated with a prediction; Bein et al., 2023). This emphasises the mixed nature of the current literature and the importance of further research into the effect of prediction on memory. If predictive processing regulates information uptake, the predictive model applied, and the predictability of the sensory information, may influence what is subsequently encoded into memory. Importantly, whilst most studies rely on changes in probability or consistency to inform the level of prediction error that is evoked (e.g., Greve et al., 2017; Ortiz-Tudela et al., 2023; Turan et al., 2023), an investigation into the neural correlates related to prediction may shed further light on the effect of predictability on memory.

### 1.3. Predictability and the N400 event-related potential (ERP)

A neural correlate frequently related to predictive processing is the N400 event-related potential (ERP), a negative-going component arising approximately 400ms post-stimulus onset (e.g., Bornkessel-Schlesewsky & Schlesewsky, 2019; Hodapp & Rabovsky, 2021; Kutas & Hillyard, 1980). The N400 exhibits greater amplitudes following incongruent or unexpected words (e.g., Federmeier & Kutas, 1999; Kutas & Hillyard, 1980, 1984), demonstrating a graded relationship with word predictability (e.g., DeLong et al., 2005; Federmeier et al., 2007; Wlotko & Federmeier, 2012). Consequently, current perspectives posit that the component reflects the degree of mismatch between a prediction and the sensory input (i.e., a prediction error; e.g., Bornkessel-Schlesewsky & Schlesewsky, 2019; Hodapp & Rabovsky, 2021). This is broadly compatible with computational modelling work (Eddine et al., 2024; Fitz & Chang, 2019; Rabovsky et al., 2018; Rabovsky & McRae, 2014, for a review see Eddine et al., 2022), highlighting the relevance of the N400 for gauging predictive processing. Although the N400 has primarily been studied in linguistic contexts, N400 effects have also been observed in non-linguistic settings (e.g., relating to action and arithmetic), suggesting that the component may embody more general functions related to meaning processing (Amoruso et al., 2013; Kutas and Federmeier, 2011; Niedeggen et al., 1999; Reid & Striano, 2008; Sitnikova et al., 2008). For example, Urgen and colleagues (2018) observed an N400 effect when participants viewed a realistic robot moving mechanically, suggesting that movement violations of appearance-based predictions may play an important role in the uncanny valley effect. However, despite the component’s applicability to non-linguistic contexts, most research relating the N400 to memory outcomes pertains to language, revealing enhanced implicit memory (as evidenced by faster reaction times) for words that evoked greater N400 amplitudes (Hodapp & Rabovsky, 2021). Meyer and colleagues (2007) also observed a correlation between N400 amplitudes to words during learning and ERP old/new effects reflecting familiarity at test, suggesting that the N400 may support familiarity-based recognition. Other research on repetition suppression provides indirect evidence supporting the role of the N400 in memory encoding, demonstrating N400 amplitude decreases following repeated exposure to incongruent sentence endings (Besson et al., 1992). This decrease in N400 activity has been interpreted to reflect adaptation of a predictive model to initially unexpected stimuli (Hodapp & Rabovsky, 2021, 2024); for similar research on the mismatch negativity (MMN) ERP, see Garrido and colleagues (2009). External to traditional language experiments, Abla and colleagues (2008) measured the N400 during a statistical learning task involving triplets of tones. Those who demonstrated stronger familiarity with the tone triplets at test showed increased N400 amplitudes for the first tone in the sequence during learning. Abla and colleagues (2008) linked this to a novelty effect due to the first tone being the least predictable. Whilst this study measured memory in the form of statistical learning, further research is necessary to test the assumption that the N400 influences behavioural memory outcomes in contexts extending beyond language.

### 1.4. The role of individual neural variability

If memory encoding is influenced by predictability, it may also be susceptible to individual differences in predictive processing. The N400 response has been shown to differ between individuals according to intrinsic neural factors, namely the aperiodic slope of the 1*/f* distribution and individual alpha frequency (IAF), the peak frequency within the alpha rhythm (Bornkessel-Schlesewsky et al., 2022; Dave et al., 2018; Jano et al., 2024; Klimesch, 1999). This may suggest that IAF and the 1*/f* slope influence predictive processing and consequent memory performance. However, to our knowledge, N400 patterns during learning have not been related to explicit memory outcomes in non-linguistic settings, and with a consideration of intrinsic neural differences. Further, while many studies examine the effect of prediction on memory with pairs or triplet relationships (e.g., Long et al., 2016; Otsuka & Saiki, 2016; Rouhani et al., 2020; Sherman et al., 2022; Sherman & Turk-Browne, 2020; Turan et al., 2023), it is important to explore this connection across longer sequences of information. The temporal accuracy of ERPs may provide a means to track predictive processing in real-time and across longer associations, to understand how predictions adapt according to information accrual. Therefore, an investigation into the relationship between the N400 and memory across sequences may have important implications for learning in naturalistic settings, where information is received in a continuous stream (Parr et al., 2022). Such an investigation may aid in understanding the consequences of predictability for learning more generally, whilst providing insight into prediction as a probabilistic and adaptive process.

### 1.5. The present study

The present study aimed to determine the effect of predictability on memory and the role of individual neural variability in this relationship. Participants first engaged in a learning task where they were exposed to predictable and unpredictable sequences of outdoor scene images, after which their recognition of the images was tested. As prior research suggests that memory is impaired for stimuli that promote predictions (Sherman & Turk-Browne, 2020), it was hypothesised that memory performance would be better for images in the unpredictable sequences, in contrast to images in the predictable sequences. Based on research highlighting the relationship between the N400 component, stimulus predictability, and memory updating (described above), it was also hypothesized that greater N400 amplitudes during learning would be associated with enhanced memory performance at recognition.

## 2. Materials and methods

### 2.1. Participants

Fifty individuals took part in the experiment; a sample size based on similar research (Sherman & Turk-Browne, 2020) that aimed to increase statistical power whilst accounting for participant drop-out rates or data corruption. Two participants were removed from the sample, one due to technical difficulties and the other due to misinterpretation of the task instructions. Consequently, the final sample size was 48 (mean age = 26.8 years old, SD = 6.6 years), consisting of 15 males and 33 females. All participants were right-handed adults between the ages of 18 and 39, with normal to corrected-to-normal vision. Participants were not taking regular medication that could impact the EEG and did not have any diagnosed psychiatric conditions (e.g., depression), diagnosed intellectual impairments, or a diagnosis of dyslexia. They also reported no recreational drug use in the past 6 months. Due to the within-groups nature of the study design, all participants were exposed to all experimental conditions (predictable and unpredictable sequences).

### 2.2. Stimuli and task

#### 2.2.1. Outdoor scene task

Stimuli during the learning phase of the outdoor scene task consisted of 102 sequences containing four outdoor scene images each (408 images total). Images were obtained from Sherman and Turk-Browne (2020), and belonged to one of twelve categories (beaches, bridges, deserts, forests, fields, lighthouses, mountains, parks, waterfalls, canyons, lakes, and reefs). In contrast to Sherman and Turk-Browne’s (2020) experiment, we included the ‘reefs’ category to replace the category ‘marshes,’ as many of the images within the ‘marshes’ category were either of a low resolution or were highly similar to each other, and we believed that they may have been too difficult to distinguish from one another. Images belonging to the ‘reefs’ category were taken from Google image searches and were identical in size to the images from the other categories (800 x 600 pixels). Images were presented in the centre of the screen using the *Opensesame* program (Mathôt et al., 2012).

Of the 102 sequences in the learning phase, 68 were predictable, as they contained the same image categories in the same A→B→C→D order (either PARK → BRIDGE → LIGHTHOUSE → WATERFALL or DESERT → BEACH → FOREST → MOUNTAIN).

Although the categories and their order remained stable within the predictable sequences, the specific image belonging to that category always differed, such that no one image was repeated during learning. This is in line with Sherman and Turk-Browne’s (2020) study, which probed item-specific memory whilst retaining predictable relationships in the form of categorical associations. The remaining 34 sequences always consisted of images from the categories CANYONS, REEFS, FIELDS, and LAKES. Images in these sequences were presented in random order, and as such, multiple images belonging to the same category could be present in the same sequence. This rendered the sequences and the images completely unpredictable, and they were hence labelled ‘Z’ images. The initial assignment of categories to the two conditions (predictable ABCD sequences and unpredictable Z sequences) was random but did not differ between participants. Further, the order in which each sequence appeared was randomised across participants, although the specific images in each sequence remained consistent between individuals. As the images belonging to 6 of the 12 overall image categories contained a man-made object, it was ensured that this distribution was balanced between the two sequence conditions, such that the predictable (ABCD) sequences contained four (out of eight total) categories with man-made objects, and the unpredictable (Z) sequences (of which there were half as many as the predictable sequences) contained two (out of four) categories with man-made objects.

During the recognition phase, stimuli consisted of the 408 ‘old’ images shown during the learning phase, belonging to both the predictable (ABCD) and unpredictable (Z) sequences. Additionally, 72 ‘new’ lure images that participants were not previously exposed to were included; 48 belonging to the predictable (ABCD) categories, and 24 belonging to the unpredictable (Z) categories. The differing number of lures between the two sequence conditions was proportional to the distribution of sequence types during learning, such that there were double the number of predictable images as compared to unpredictable images (note that this was considered when calculating memory performance). To maintain consistency regarding the distributions of the various image categories, six lures belonged to each of the 12 outdoor scene categories, resulting in 12 lure images per A, B, C, D position for the predictable ABCD sequences. For example, as A images belonged to the ‘park’ and ‘desert’ categories, six lures were images of parks, and six were images of deserts. As the Z condition consisted of four image categories (CANYONS, REEFS, FIELDS, and LAKES), six lures from each category were presented. The number of lures per category is adapted from Sherman & Turk-Browne (2020), who presented participants with predictable A → B pairings and included four lures per predictable (AB) category in their recognition phase. As our design was extended to include longer image sequences and unpredictable sequences, we chose six lures per category to account for this increase. All images during the recognition phase were presented sequentially and in a random order across participants. A schematic of the outdoor scene task is presented in *Figure 1*.

**Figure 1.**
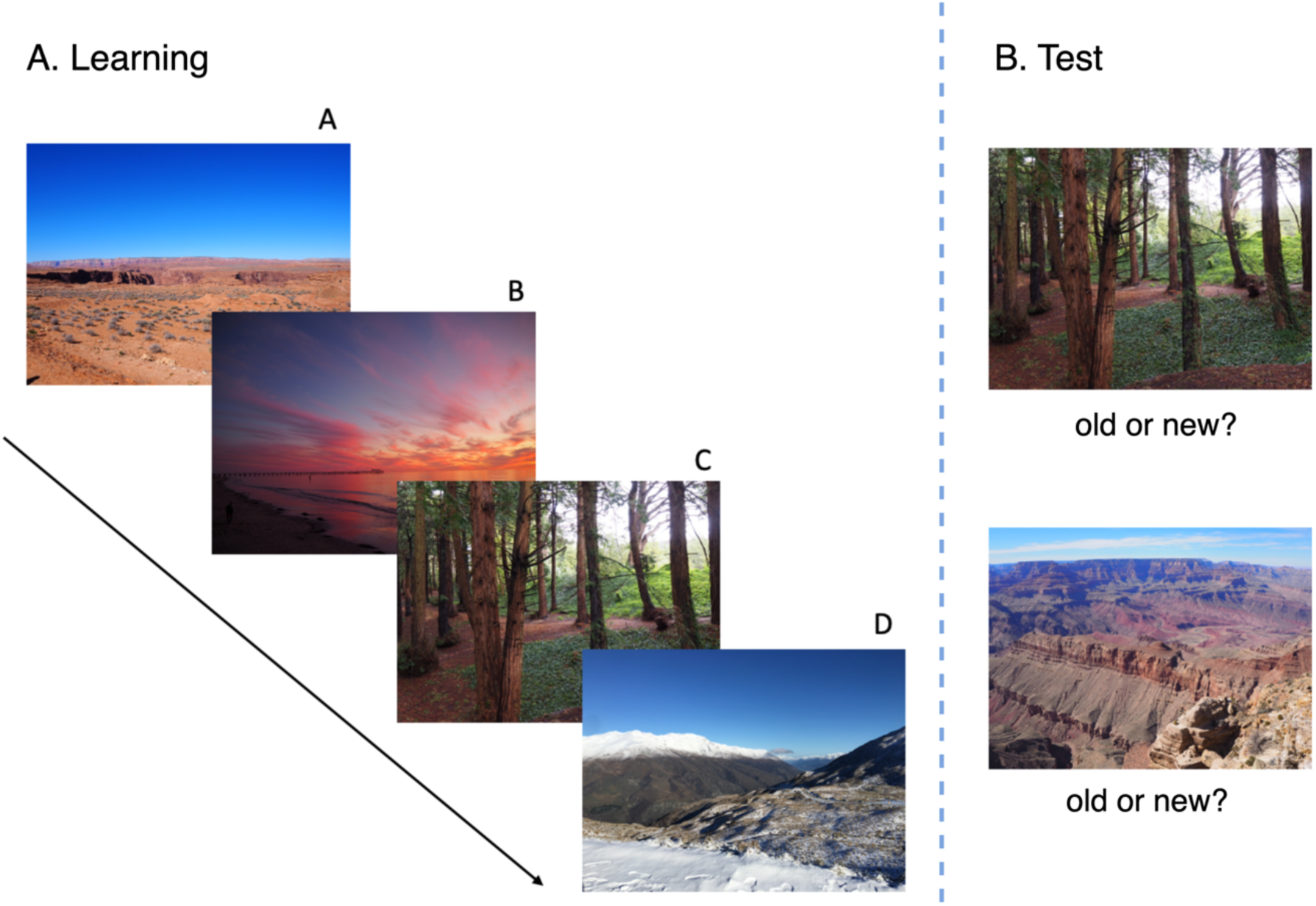
Schematic of the outdoor scene task. **(A)** The learning phase of the outdoor scene task, where participants were presented with predictable and unpredictable four-item sequences of outdoor scene images. Each image was presented sequentially and was separated by a fixation dot. This example is of a predictable ABCD sequence (A: desert, B: beach, C: forest, D: mountain). **(B)** The recognition test phase of the outdoor scene task, where participants were presented with old images from the original learning phase, and new lure images, to which they made an ‘old’ or ‘new’ judgement.

#### 2.2.2. Pilot testing

Prior to the main study, a brief pilot test with five participants who did not take part in the main experiment was conducted to determine whether individuals were able to learn the predictable four-item ABCD sequences. The same predictable sequences as in the main experiment were used, and participants completed the learning phase as usual. However, instead of a following recognition test, participants engaged in a simple statistical learning test, similar to the pattern completion tests reported in Siegelman et al. (2017). Participants viewed the A, B, and C images in the original predictable sequences. However, for the final D image, they were required to choose between two images; the image that was present in the original sequence, and a foil image belonging to a different image category. Feedback was not provided throughout the task. Mean accuracy on this task was 74%, suggesting that the design facilitated sequence learning.

### 2.3. EEG measures

Electroencephalography (EEG) was recorded throughout the experimental tasks using a 32- channel Brainvision ActiCHamp system (Brain Products GmbH, Gilching, Germany). For the reference electrode, the T7 electrode in the Afz position was used. However, one participant had a faulty T7 electrode, and so the FT9 electrode was used as the reference instead. The ground electrode was placed in the Fpz position and dropdown electrodes from the Fp1 and Fp2 positions were used as the bipolar electrooculogram (EOG) channels, placed above and below the left eye. The data were sampled at a frequency of 500Hz (DC coupled), and the impedances of the channels were maintained below 10kΩ.

### 2.4. Experimental design and procedure

Ethical approval for the study was first sought from the University of South Australia Human Research Ethics committee. Eligible participants provided written informed consent to participate in the research, after which their EEG was set up by an experienced technician. They were seated at a computer in a quiet testing room, with their eyes closed for two minutes, and then with their eyes open for two minutes, to obtain recordings of their resting state brain activity for the calculation of IAF and aperiodic activity. Following this, participants either engaged in the main outdoor scene task, or completed short control tasks. The order in which participants completed the main task and the control tasks was counterbalanced between participants to control for fatigue effects.

The procedure for the outdoor scene task is loosely based on Sherman & Turk-Browne (2020)’s protocol. For the learning phase of the outdoor scene task, participants viewed the four-item sequences whilst their EEG was recorded. They were not informed of the underlying patterns amongst the images but were simply told that each sequence would consist of four images, presented one after the other, to which they must pay attention. Here, we avoided specifically instructing participants to predict the upcoming items, as prior research suggests that predictive mechanisms are engaged in the presence of probabilistic regularities (Baker et al., 2014; Bar, 2007; Dale et al., 2012; Mumford, 1992). Additionally, while some studies encourage participants to actively predict (e.g., Ortiz-Tudela et al., 2023; Turan et al., 2023), if prediction is a general mechanism underlying cognition (Huang & Rao, 2011), doing so may engender a form of explicit anticipation that differs from naturalistic predictive processing. Each image was presented for 1000ms, followed by a fixation dot in the centre of the screen that was present for 500ms. The between-sequence intervals were 2000ms in duration, to create noticeable boundaries between sequences. As a secondary cover task, throughout the learning phase, a prompt would appear on the screen telling participants to press ‘Enter,’ as quickly as possible. This would occur at seven random between-sequence intervals during the experiment and would time-out only once the enter key had been pressed. Participants were informed that this was a secondary task to ensure that they were paying attention. The total duration of the learning phase was approximately 15 minutes.

After the learning phase, participants performed a digit span task to minimize recency effects. The task was presented via the OpenSesame program (Mathôt et al., 2012) and took approximately 15 minutes to complete. Following this, participants engaged in a surprise recognition test, where they were presented with old images and new lure images in a random order whilst their brain activity was recorded. For each image, participants were required to state whether the image was ‘old’ (they saw it in the original learning phase), or ‘new’ (they did not remember seeing it during the learning phase), using the ‘z’ and ‘m’ keys, respectively. Participants responded to the image whilst it was on the screen, with a keypress triggering a switch to a fixation dot with a duration of 500ms. Image presentations timed out after 6 seconds if no response was provided. The duration of the recognition phase was approximately 15 minutes.

The control tasks, completed either before or after the outdoor scene task, consisted of a visual statistical learning (VSL) task (based on Siegelman et al., 2017) that took approximately 20 minutes to complete, and other short tasks. As the control tasks are not the focus of the current investigation, they will not be discussed further. Following completion of the experiment, the EEG cap was removed, and participants were provided with a $50 payment as compensation for their time. The total testing time for the experiment was approximately 2 hours.

## 3. Data analysis

### 3.1. d-prime (d’) calculation

To quantify participants’ recognition accuracy, d-prime (d’; McNicol, 2004) was calculated based on performance during the recognition testing phase of the outdoor scene task. Performance was grouped according to participants’ old/new responses, where the number of hits (old images correctly identified as ‘old’) and false alarms (new images incorrectly judged as ‘old’) were calculated per image position/condition (predictable A, B, C, D, and unpredictable Z), across participants. Following this, d’ scores for each image type (predictable ‘A’, ‘B’, ‘C’, ‘D’ and unpredictable ‘Z’ images) were computed using the *psycho* package in R version 4.3.1, which subtracts the z-score of the false alarm rate from the z-score of the hit rate (Makowski, 2018). Six subjects with false alarm rates equal to zero were identified, and as such, the d’ function was run with adjust = ‘TRUE’ to adjust for extreme scores in line with Hautus (1995). Note that it was not possible to obtain d’ scores for each image position for images in the unpredictable ‘Z’ condition. This is because lure images were based on categories, but the categories’ positions in the unpredictable sequences differed each time (e.g., for one sequence, an image of a canyon might be in the ‘A’ position, but for the next sequence, a canyon image might be in the ‘C’ position). As such, d’ for unpredictable ‘Z’ images was calculated across all image positions.

Unforeseen technical difficulties were experienced by five participants during the learning phase of the outdoor scene task, such that the task crashed at variable positions. For one participant, the task crashed towards the beginning (after the 32^nd^ sequence), and they completed the learning phase again. The trials that they had seen twice were removed from their behavioural recognition results in case repeated exposure led to enhanced memory performance for those items. For the remaining four participants, the trials that they had not seen during learning were removed from their behavioural recognition results. We adjusted the d’ calculations for all of these participants based on the number of lure and target images that were seen. However, one participant appeared to have made an abnormally large number of ‘new’ judgements, especially to ‘old’ images that were seen during learning. As this may have reflected a bias in performance (potentially resulting from the participant’s belief that they had not seen many of the images), this participant was removed from subsequent analyses.

### 3.2. EEG data processing

To maintain EEG files in line with recommended standards, the EEG data from the learning phase were first transformed into the Brain Imaging Data Structure (BIDS; Appelhoff et al., 2019) using the *MNE-BIDS* package in MNE-Python (Gramfort et al., 2013; note that Python version 3.8.8 was used). Subsequent pre-processing occurred using MNE-Python, where the data were re-referenced to the mastoid channels (TP9 and TP10). Power spectral density (PSD) plots were then generated, and the data were visually inspected for bad channels, which were marked. EOG channels were set as Fp1 and Fp2 before an independent component analysis (ICA) was fit to a copy of the data that was band-pass filtered between 1 to 40Hz. The ICA was then applied to the raw data to identify and remove ocular noise, after which bad channels were interpolated using spherical splines. Next, the data was filtered with a bandpass filter ranging from 0.1 to 30Hz.

Following processing, the data were epoched around events of interest, between -0.3ms preceding the images to 1 second post-image (1.3 seconds total duration). The *Autoreject* function (Jas et al., 2016) was then applied to interpolate and/or remove bad epochs, resulting in an average of 246.9 epochs for predictable images and 123.2 epochs for unpredictable images across participants.

### 3.3. ERP generation

Event-related potentials (ERPs) were computed using MNE-Python (Gramfort et al., 2013) over an N400 window that ranged from 300ms to 500ms post-image onset, based on prior literature (Federmeier et al., 2007; Hodapp & Rabovsky, 2021; Hubbard et al., 2019; van Berkum et al., 2003). The mean N400 amplitude within the window of interest, per channel, trial and participant was calculated. For a discussion of the chosen N400 topography, see *Results* section.

Rather than performing a traditional baseline correction, ERP activity was additionally computed over a pre-stimulus window ranging from -300 to 0ms prior to image presentation. Mean pre-stimulus ERP amplitudes were calculated for each channel and trial, and were used as a ‘baseline’ window, that was included as a covariate in statistical models. This approach is described by Alday (2019) and is proposed to enhance statistical power in contrast to traditional baseline correction methods. To generate ERP grand average plots, the data were resampled to 125Hz, and frequency and amplitude data at each time point was computed. Grand averages were plotted using the *ggplot2* package in R version 4.3.1 (Wickham, 2016).

### 3.4. Individual metrics

To obtain measures of individual alpha frequency (IAF) and the 1*/f* aperiodic slope, eyes closed data from resting state recordings was used. The data were re-referenced to TP9 and TP10 and filtered with a bandpass filter from 0.1 to 30Hz. IAF was calculated using the *Philistine* package in MNE-Python (Alday, 2018), over electrodes P3, P4, Pz, O1, O2 and Oz, as IAF is typically measured in parietal-occipital areas (e.g., Corcoran et al., 2018; Dziego et al., 2023; Grandy, Werkle-Bergner, Chicherio, Schmiedek, et al., 2013). The lower bound of the alpha frequency was set at 7Hz, and the upper limit was 13Hz. This method is based on Corcoran et al. (2018) and calculates both the peak alpha frequency (PAF; the most prominent peak in the alpha spectrum) and the centre of gravity (CoG; the weighted mean of power across the alpha frequency band; Corcoran et al., 2018).

To obtain estimates of the aperiodic 1/*f* slope, the power spectral density was first calculated across the raw data, for channels P3, P4, Pz, Oz, O1 and O2, to maintain consistency with the IAF metrics. The *Fitting Oscillations and One-Over f (FOOOF)* package (version 1.0.0; Donoghue et al, 2020) was then used to parameterize the power spectrum, which occurred across a 1 to 30Hz frequency range. The *FOOOF* model settings included a fixed aperiodic mode with a peak width limit of two to eight, a peak threshold of two, a minimum peak height of zero, and a maximum number of four peaks. Subsequently, the data were averaged across the three channels, resulting in a single 1/*f* aperiodic slope measure per participant.

### 3.5. Statistical analysis

Linear mixed-effects models (LMMs) were computed with the *lme4* package in R version 4.3.1 (Bates et al., 2015) to examine effects of interest, as such models allow for the specification of both fixed and random factors (See *Table 1* for statistical model structures). Whilst random intercepts were included in all models, the inclusion of by-subject and/or by-item random slopes led to convergence/singularity issues for some. As such, random slopes were omitted in some cases, as this may reflect overfitting (for a discussion of model overparameterization, see Bates et al., 2018). To calculate *p*-values, type II Wald tests from the *car* package were used (Fox & Weisberg, 2019), which were computed according to an alpha level of 0.05. The categorical variables of condition and position were sum contrast coded using the *contr.sum* function from the *car* package (Fox & Weisberg, 2019), and continuous predictors were scaled and centred. Model effects were visualised with 83% confidence intervals using *ggplot2* (Wickham, 2016). In some cases, IAF and the 1/*f* slope were included in the same model as separate interaction terms to reduce the number of models. Also note that for statistical models including IAF, CoG is specifically used as it may perform better when a discernible alpha peak is not present (e.g., in cases where a split peak is observed; Corcoran et al., 2018; Klimesch, 1997; Klimesch et al., 1993). Finally, as IAF could not be estimated for two participants, models with IAF were computed with a sample size of 46, whilst models that did not include the factor of IAF included the full sample size of 48 participants. Models were fitted using variables of interest. However, if a predictor did not interact to influence the outcome variable, and if it did not provide a grouping structure for the data, it was removed to reduce model complexity. Note that this resulted in the removal of a model predicting d’ from 1/*f* slope, condition, position and N400 amplitude, as 1/*f* slope did not influence the outcome variable. As such, the model did not contribute to the results over and above Model 5 (see *Appendix A, Model 5.2* for model output).

**Table 1.** Linear mixed-effects model structures including outcome variables, fixed effects, covariates, random intercepts, and random slopes.

| ***Behavioural models*** |  |  |  |  |  |
| --- | --- | --- | --- | --- | --- |
| **Model** | **Outcome** | **Fixed effects** | **Separate interaction terms** | **Random intercepts** | **Random slopes** |
| 1 | d' | Condition (predictable/unpredictable) | 1/f slope<br>IAF | + Subject |  |
| 2 | d' | Image type (predictable A/B/C/D, unpredictable Z) | 1/f slope<br>IAF | + Subject |  |
| ***ERP models*** |  |  |  |  |  |
| **Model** | **Outcome** | **Covariates** | **Fixed effects** | **Random intercepts** | **Random slopes** |
| 3 | N400 amplitude (300-500ms) | Pre-stimulus ERP amplitude | Sagittality (anterior/central/posterior)<br>Laterality (left/middle/right) | + Subject<br>+ Item |  |
| 4 | N400 amplitude (300-500ms; averaged over anterior channels) | Pre-stimulus ERP amplitude | Condition (predictable/unpredictable)<br>Position (A/B/C/D) | + Subject<br>+ Item |  |
| ***Combined ERP and behavioural models*** |  |  |  |  |  |
| **Model** | **Outcome** | **Covariates** | **Fixed effects** | **Random intercepts** | **Random slopes** |
| 5 | d' | Pre-stimulus ERP amplitude | Condition (predictable/unpredictable)<br>Position (A/B/C/D)<br>N400 amplitude (300-500ms; averaged over anterior channels)<br>IAF | + Subject | By-subject according to condition |

## 4. Results

Statistical model outputs are presented in Appendix A, whilst the data and analysis scripts necessary for replication of the results are freely available at the following repository: https://osf.io/qvjer/.

### 4.1. Descriptive statistics

The mean reaction time in response to the ‘enter’ prompt during the cover task of the learning phase of the outdoor scene task was 0.770 seconds (*SD* = 0.399 seconds). Mean and standard deviation d’ scores from the learning phase of the outdoor scene task are presented in *Table 2*.

**Table 2.**
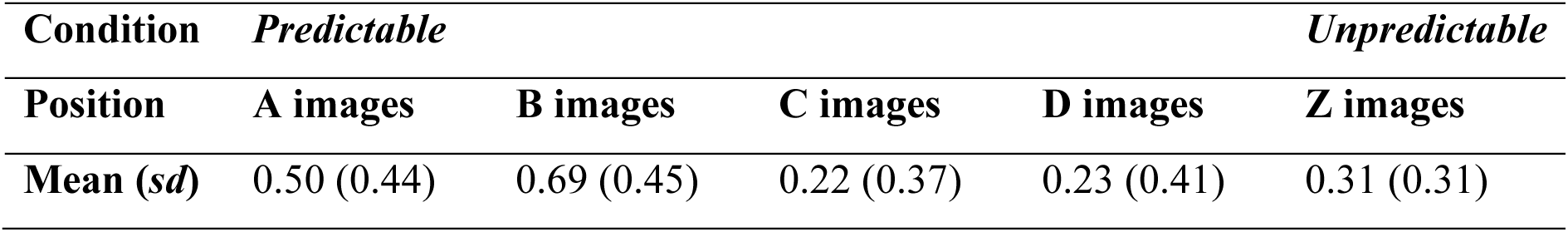
Mean d’ scores per image condition and position, with standard deviations in brackets. Note that d’ was computed across all images in the unpredictable ‘Z’ sequences, regardless of their position, as category lures could not be made specific for each position.

### 4.2. Linear mixed model results of primary analyses

#### 4.2.1. Behavioural results

To investigate the hypothesis that predictable images would be remembered more poorly than unpredictable images, Model 1 was fitted to the data including the variables of image condition (predictable/unpredictable), IAF and 1/*f* slope. The model detected a significant main effect of *condition* on d’ (χ2(1) = 4.50, *p* = .034; See *Figure 2a*). However, the interaction between *IAF* and *condition* was nonsignificant (χ2(1) = 1.54, *p* = .214), in addition to the interaction between *condition* and *1/f slope* (χ2(1) = 3.00, *p* = .083).

**Figure 2.**
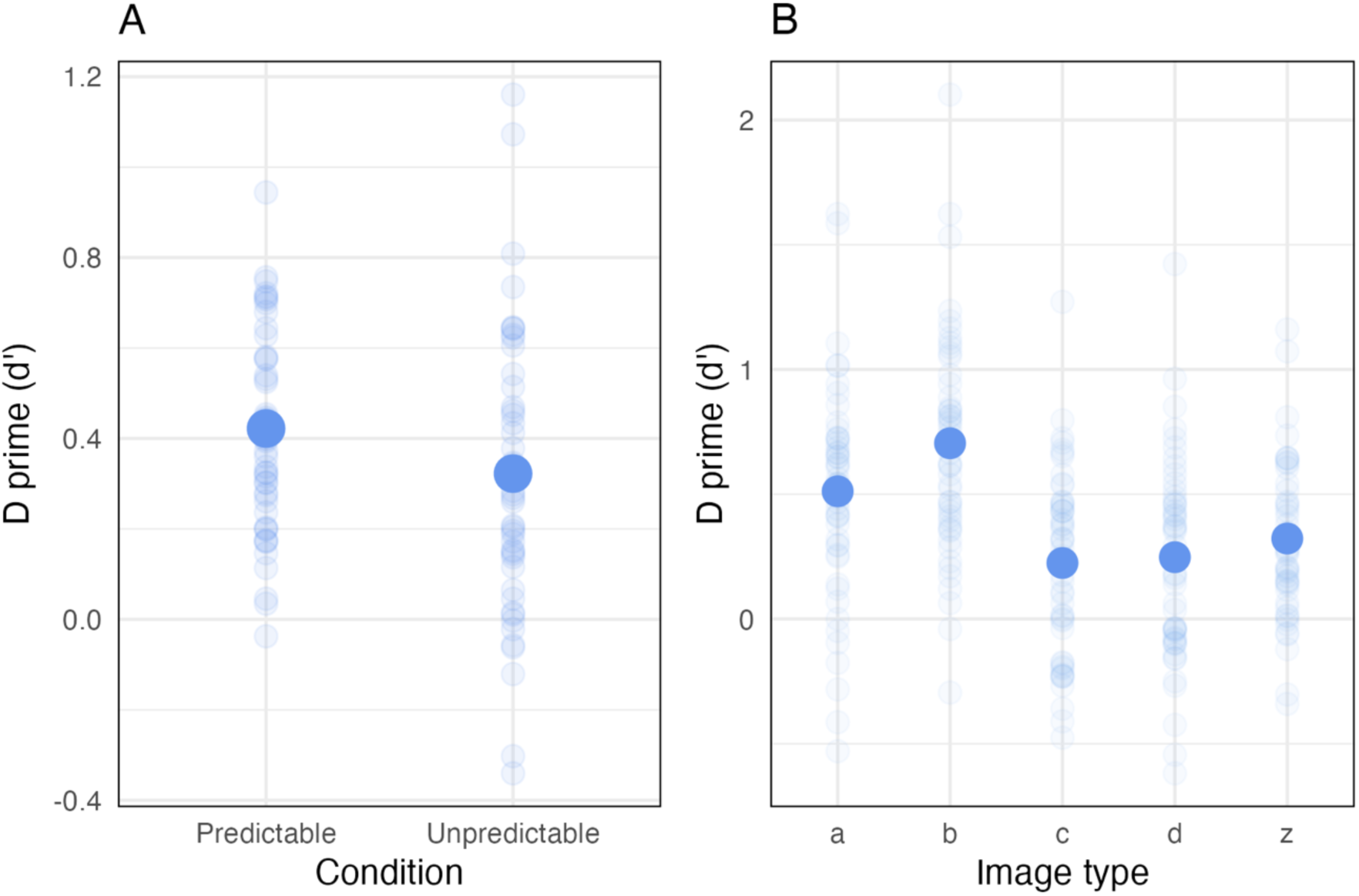
Behavioural data. **(A)** d’ score predicted by image condition (predictable versus unpredictable). **(B)** d’ score predicted by image type (predictable A, B, C, D, and unpredictable Z images). Faded dots in both plots indicate raw data points, and filled dots reflect predicted model values.

To investigate the effect of both image condition and image position, the ‘image type’ variable was used for the behavioural analyses, as the levels of this variable were divided into predictable ‘A’, ‘B’, ‘C’ and ‘D’ images, and unpredictable ‘Z’ images. To investigate the effect of image type, IAF and the 1/*f* slope on d’ scores, Model 2 was computed. A significant main effect of *image type* on d’ was detected (χ2(4) = 50.97, *p* <.001), such that d’ scores were greatest for predictable B images (See *Figure 2b*).

#### 4.2.2. N400 results

To determine a region of interest for analyses of the N400, Model 3 predicted N400 amplitude from sagitallity and laterality. A significant interaction between *sagitallity* and *laterality* was observed (χ2(4) = 175.24, *p* <.001), whereby N400 amplitude was largest over anterior regions, particularly over the midline area (for topography map and grand average across all regions, see *Figure 3*). This is consistent with previous research, which suggests that the N400 in pictorial paradigms exhibits a frontal topography (Kutas & Federmeier, 2011; Urgen et al., 2018). As such, for subsequent models including N400 amplitude, the data were filtered to capture activity over anterior electrodes (Fz, F3, F7, FT9, FC5, FC1, FT10, FC6, FC2, F4, F8). This method was chosen as the inclusion of the variables ‘sagitallity’ and ‘laterality’ in other models (which consisted of additional predictors) would otherwise likely lead to overcomplexity that would not be supported by the number of observations. Additionally, whilst we detected an interaction between both sagitallity and laterality, we opted to include all anterior electrodes, as only a single electrode was classified as belonging to the anterior-mid region (Fz).

**Figure 3.**
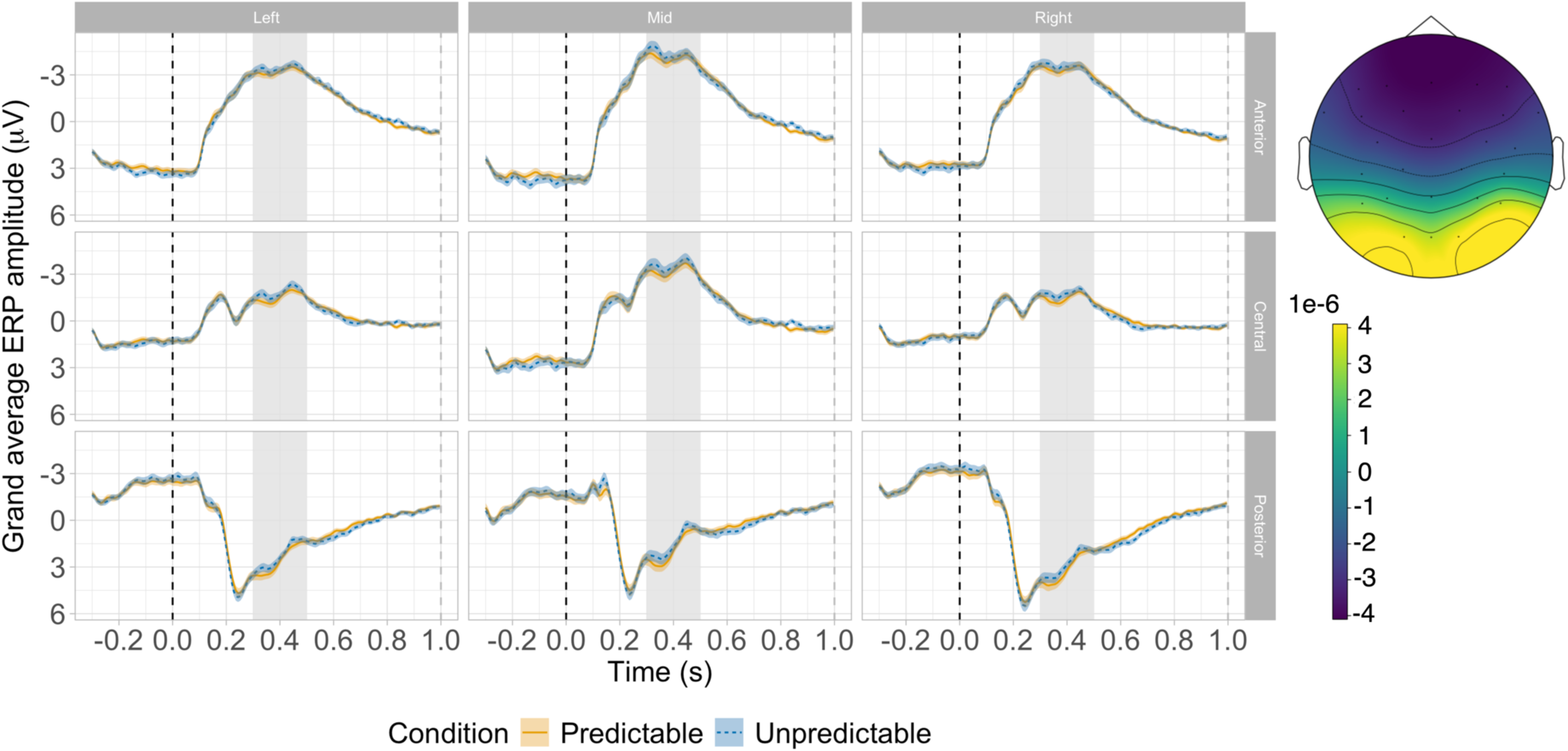
Grand average ERPs according to condition. Grand average plots depicting mean ERP activity according to image condition (predictable versus unpredictable), sagitallity (anterior, central, posterior) and laterality (left, mid, right). Black dotted lines represent image onset, grey dotted lines mark the end of image presentation, and grey shading depicts the N400 window used in statistical analyses. Orange lines reflect ERP activity in the predictable condition, whilst blue dotted lines represent the unpredictable condition. Orange and blue shading depicts the standard errors for each condition. A topography map in the bottom right-hand corner displays the mean ERP activity across the scalp in the N400 window (300 to 500ms post-stimulus onset), with the colour gradient (light to dark) representing ERP amplitude in volts.

Model 4 was fitted to examine the effects of image condition and position on the amplitude of the N400 component (averaged over anterior channels to reduce complexity). Note that as ERPs were computed at the single trial level (versus subject level for d’ scores), the variables ‘condition’ and ‘position’ contained differing information and could thus be included in the same model, in contrast to the behavioural models, where ‘image type’ is used. While the interaction between image *condition* and *position* was non-significant (χ2(3) = 3.02, *p* = .388), the main effect of *position* was significant (χ2(3) = 21.28, *p* <.001). N400 amplitudes were greatest for images in the ‘B’ position (see *Figure 4 and Figure 5A*).

**Figure 4.**
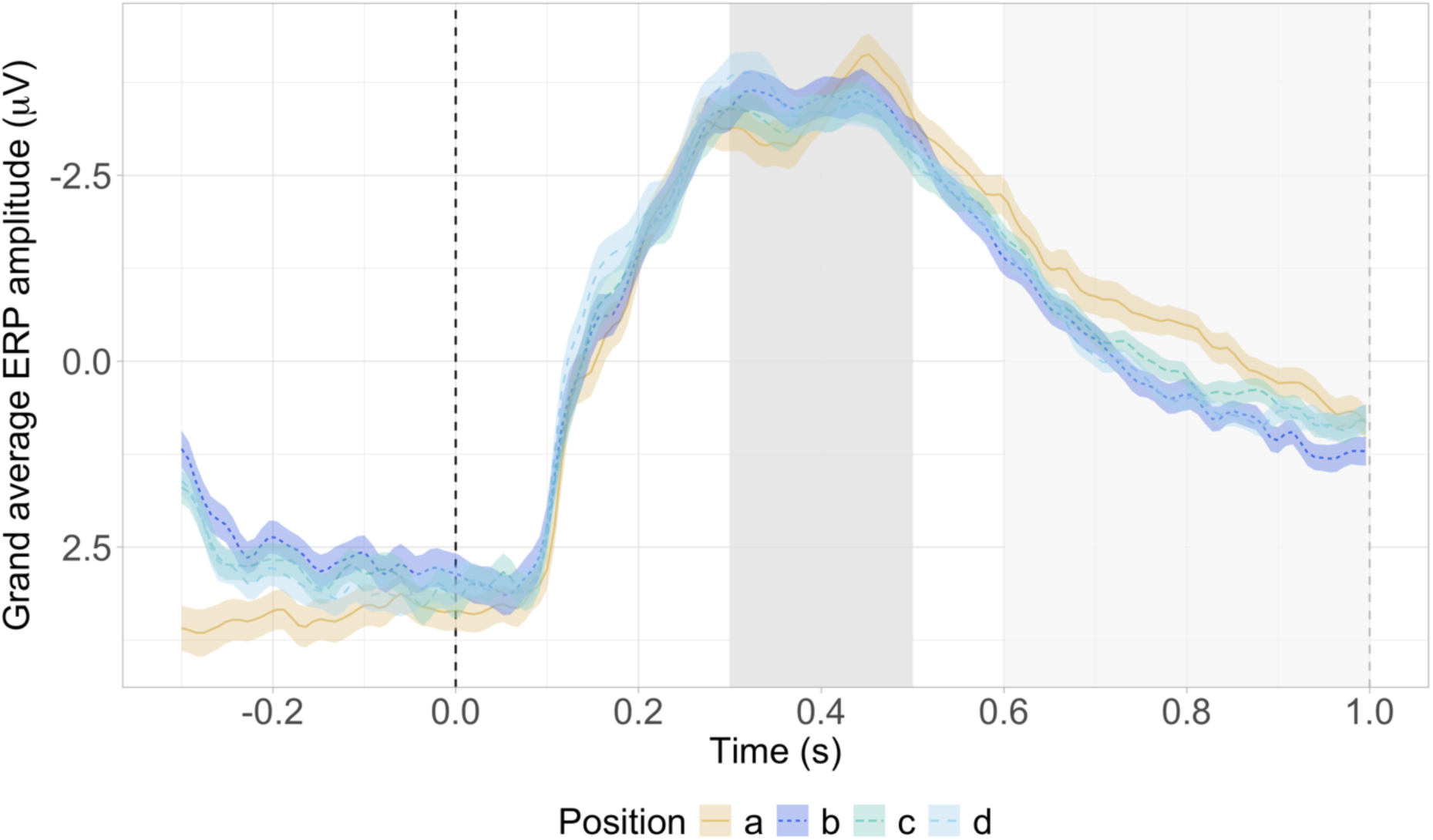
Grand average ERP activity according to position. Grand average plots depicting mean ERP activity according to image position across all anterior channels. Black dotted lines represent image onset, grey dotted lines mark the end of image presentation. The first, darker grey box depicts the N400 window used in statistical analyses, while the second, lighter grey box shows the time window used for exploratory statistical analyses (described in section 4.3).

**Figure 5.**
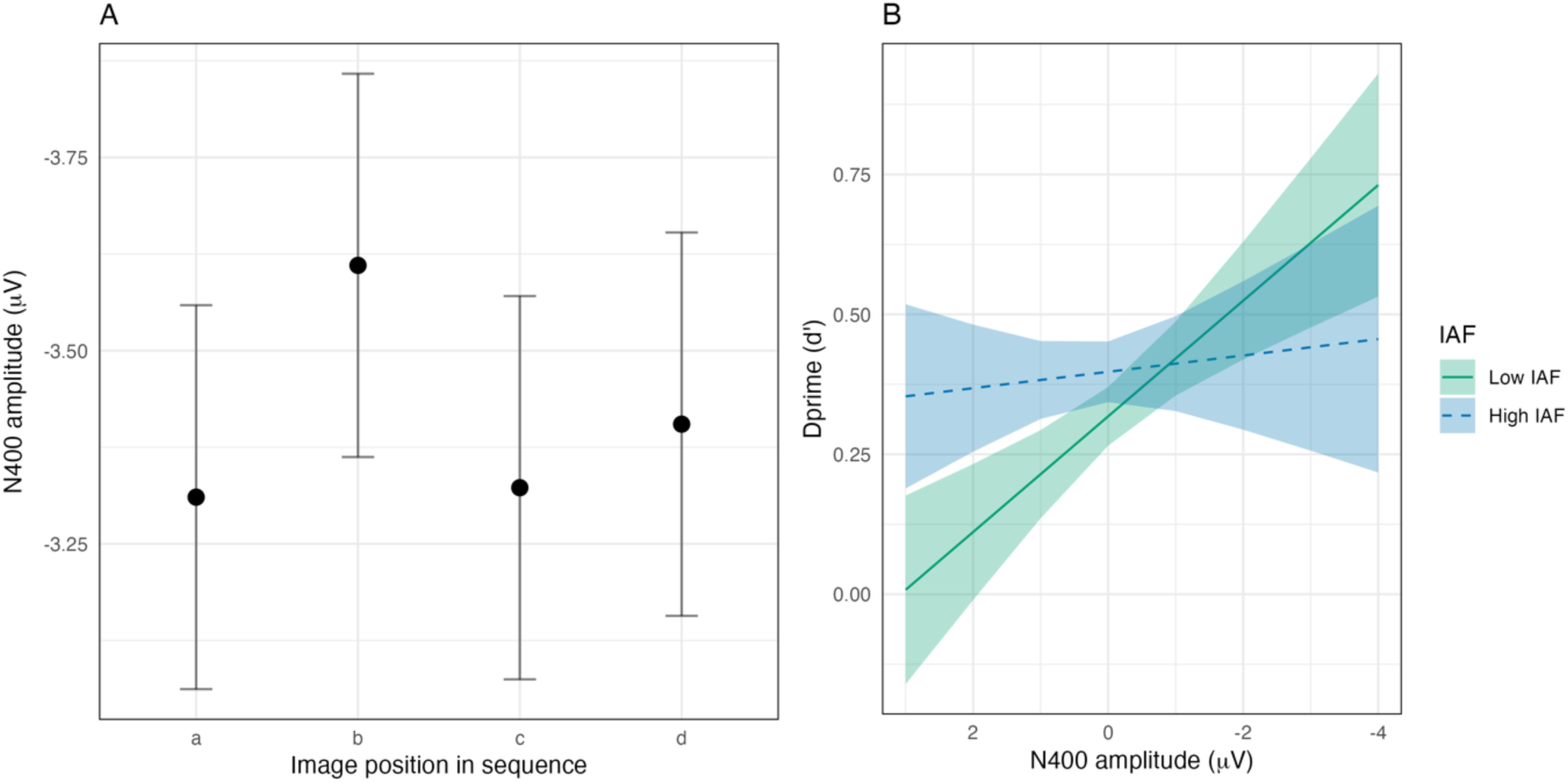
N400 effects. **(A)** The modelled effect of image position (A, B, C, D) on N400 amplitude. Error bars reflect 83% confidence intervals. **(B)** Linear mixed model results depicting the interaction between N400 amplitude and IAF on d’. Green lines represent low IAFs, whilst blue dotted lines depict high IAFs, with this dichotomisation calculated according to the first and third quartiles for visualisation purposes only. Green and blue shading depicts the 83% confidence intervals.

#### 4.2.3. Combined N400 and behavioural results

It was predicted that images that evoked greater N400 amplitudes during learning would be better remembered. To investigate this hypothesis, we ran a combined LMM (Model 5) predicting d’ from image condition (predictable/unpredictable), position (A, B, C, D), N400 amplitude, and IAF. Importantly, as d’ is an aggregated variable, this analysis could not be completed at the single trial level. Instead, N400 amplitude was averaged such that a mean N400 amplitude score per image condition, position, and subject, was computed. A significant interaction between *N400 amplitude* and *IAF* was revealed (χ2(1) = 7.59, *p* = .006), such that for low IAF individuals, d’ scores increased as N400 amplitudes during learning increased. This pattern was not observed for those with a high IAF (See *Figure 5B*).

### 4.3. Exploratory late positivity models

ERP grand average plots (specifically *Figure 4* according to image position) revealed differences at a later time window resembling a late positive complex (LPC) ERP. As an additional exploratory analysis, we computed this component over a 600ms to 1000ms window post-image onset. Prior research generally examines the LPC over an approximate 500ms to 800ms window post stimulus onset (Stróżak et al., 2016; Yang et al., 2019), although it has been measured within a 500ms to 1000ms window before (Brezis et al., 2017). We extended the typical window to capture the prominent differences in the component (as identified via visual inspection), whilst avoiding overlap with the subsequent inter-stimulus interval. A data-driven approach was taken to determine the region of interest for this component, similarly to the N400 models, where sagitallity and laterality were first entered into a model. Following this, the region demonstrating the largest amplitude was used for subsequent models (for exploratory model structures see *Table 3*).

**Table 3.** Linear mixed-effects model structures for exploratory models including outcome variables, fixed effects, covariates, random intercepts, and random slopes.

| *ERP models* |  |  |  |  |  |
| --- | --- | --- | --- | --- | --- |
| Model | Outcome | Covariates | Fixed effects | Random intercepts | Random slopes |
| 6 | Amplitude of late positivity | Pre-stimulus ERP amplitude | Sagittality (anterior/central/posterior)<br>Laterality (left/middle/right) | + Subject<br>+ Item |  |
| 7 | Amplitude of late positivity (averaged over anterior channels) | Pre-stimulus ERP amplitude | Condition (predictable/unpredictable)<br>Position (A/B/C/D) | + Subject<br>+ Item | By-subject according to condition |
| *Combined ERP and behavioural models* |  |  |  |  |  |
| Model | Outcome | Covariates | Fixed effects | Random intercepts | Random slopes |
| 8 | d' | Pre-stimulus ERP amplitude | Condition (predictable/unpredictable)<br>Position (A/B/C/D)<br>amplitude of late positivity (averaged over anterior channels) | + Subject | By-subject according to condition |

#### 4.3.1. Late positivity results

Model 6 tested the effect of sagitallity and laterality on the amplitude of the late positive component, revealing a significant interaction between *sagitallity* and *laterality* (χ2(4) = 156.53, *p* <.001). Amplitude was greatest over anterior midline regions. However, similarly to the N400, all anterior channels were chosen for subsequent analyses as only a single electrode belonged to the anterior midline region (Fz).

Model 7 examined the effect of image condition and position on late positivity amplitude (averaged across anterior channels). The effect of *condition* and *position* on the amplitude of this later component was significant (χ2(3) = 8.32, *p* = .040), such that amplitudes were greatest for unpredictable versus predictable images at the ‘A’ position, whilst at the ‘D’ position, this pattern reversed (See *Figure 6A*).

**Figure 6.**
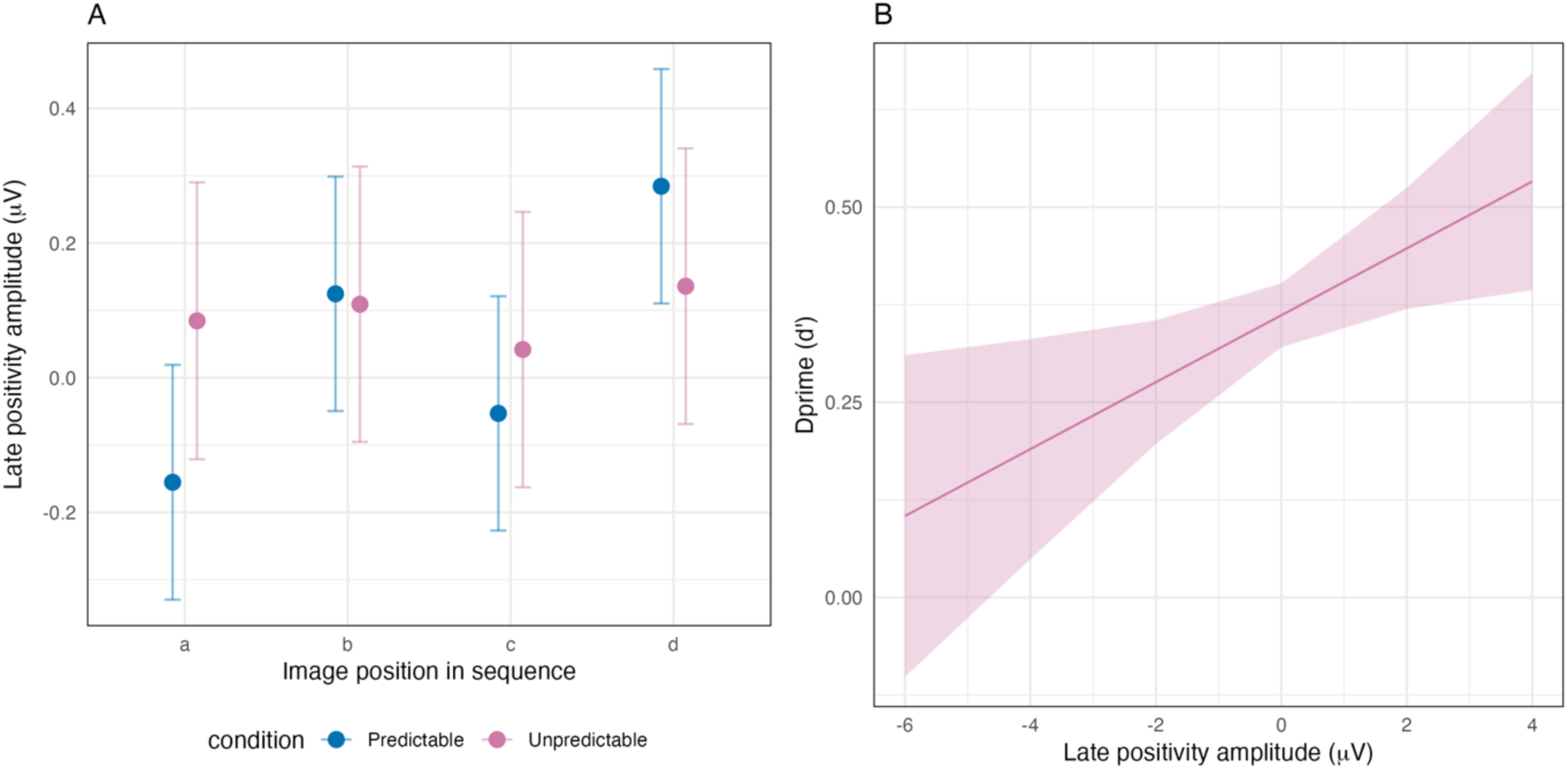
Late ERP effects. **(A)** The modelled effect of image position (A, B, C, D) and condition (predictable, unpredictable) on the amplitude of the late positivity. Error bars reflect 83% confidence intervals. **(B)** Linear mixed model results depicting the effect of late positivity amplitude on d’. Pink shading depicts the 83% confidence interval.

#### 4.3.2. Combined behavioural and late positivity results

Finally, the effect of late positivity amplitude, condition and position on d’ was investigated via Model 8. While the interaction of all three variables was non-significant (χ2(3) = 7.48, *p* = .058), the main effect of *late positivity amplitude* on d’ was significant (χ2(1) = 4.07, *p* = .044), such that the two variables were positively correlated (See *Figure 6B*). As such, the amplitude of this later component differed according to image position and condition and was predictive of later memory performance.

## 5. Discussion

The present experiment aimed to shed light on the relationship between predictability and memory, by relating ERP activity during exposure to predictable and unpredictable sequences to memory recognition outcomes. Contrary to hypothesised effects, the results revealed enhanced memory for images in predictable, as compared to unpredictable conditions. However, when taking image position into account, memory performance was best for predictable images in the ‘B’ sequence position. This finding will be subsequently discussed in relation to precision and attentional mechanisms. Strikingly, greater N400 amplitudes during learning were associated with better memory outcomes at test for low versus high IAF individuals. This may imply that intrinsic neural differences modulate the extent to which N400 activity influences memory processing. Note however, that the 1/*f* aperiodic slope did not affect this relationship, suggesting that IAF may be particularly relevant for the association between neural activity and subsequent memory. Additionally, exploratory analyses revealed changes in a later positive component (potentially resembling the LPC) according to image position and condition. This component was positively associated with later memory performance, emphasising the relevance of this activity for learning. Overall, the current findings shed new light on the interconnectedness between predictability and memory, whilst emphasising the importance of between-subject neural variability for memory encoding.

### 5.1. Changes in precision may influence the extent of memory encoding

Memory recognition was strongest for images in the predictable versus unpredictable condition, calling to question the proposed trade-off between prediction and memory encoding (Hubbard et al., 2019; Sherman et al., 2022; Sherman & Turk-Browne, 2020). However, while at a surface level, this finding may suggest that predictability enhances memory, the observation that predictable ‘B’ images were best remembered could highlight a role for attentional mechanisms in this relationship. Not only was the ‘B’ image predictable and predictive of the upcoming ‘C’ and ‘D’ items, but it was likely the first image that confirmed whether the sequence was unfolding in a predictable manner. As such, it is possible that predictable images in this position were attended to more strongly than other images, due to their relevance for evaluating the sequence. In predictive coding, attention is proposed to reflect judgements concerning the *precision* of a stimulus; that is, the amount of certainty or variance in the signal (Bastos et al., 2012; Feldman & Friston, 2010; Friston et al., 2011). Increases in attention may be associated with increases in the precision of sensory signals, enhancing the extent to which the stimulus prompts model updating (Hohwy, 2012). In the present study, attention might have been directed towards the predictable ‘B’ images once participants learnt their relevance for the broader sequence, leading to increased precision-weightings. Stronger precision-weightings could have consequently triggered a switch to memory encoding, thus explaining the enhanced memory for the ‘B’ image.

This explanation is further complemented by the observation that N400 amplitudes were generally greatest for images in the ‘B’ position, regardless of image condition (predictable/unpredictable). The N400 is proposed to not only reflect a prediction error, but to be indicative of a *precision-weighted* prediction error signal that is dependent on stimulus uncertainty (Bornkessel-Schlesewsky & Schlesewsky, 2019). As such, greater N400 amplitudes may reflect stronger precision-weightings, potentially due to enhanced attention toward the ‘B’ images. These results therefore suggest that in longer sequences, the brain may prioritise the items deemed most informative, and direct attention away from subsequent items if the prior information is sufficient for predictive processing. This is supported by Darriba & Waszak (2018), who investigated predictions across four-item sequences. Increases in the P2a and P3 ERP components (proposed to reflect evidence accumulation) were observed from the first to the third stimulus, but not at the final stimulus in the predictable sequences (Darriba & Waszak, 2018). Consequently, it was suggested that evidence accumulates until sufficient information for the prediction is acquired, potentially explaining the prioritisation of the ‘B’ image observed here. However, it is important to note that whilst we assume that the ‘B’ images received more attention, this was not explicitly measured, and future research may benefit from relating more direct measures of attention (e.g., eye-tracking; Carter & Luke, 2020) to prediction during sequence learning.

To explain why the above N400 effects occurred regardless of image condition, we also highlight a crucial aspect of our study design: whilst predictable sequences promoted predictions about upcoming image *categories*, it is unlikely that *item-specific* predictions could be precise enough to lead to N400 reductions, because each item was unique. This distinction between category and item-based predictions is congruent with hierarchical predictive coding views, where predictions at higher levels may code for more complex information (e.g., sequential categorical associations), whilst predictions at lower levels pertain to more basic (e.g., item-specific) attributes (Rao & Ballard, 1999; Summerfield & DeLange, 2014). As such, memory for images in predictable sequences might have been enhanced at the item-level, which importantly, is also the level at which memory recognition was tested. Therefore, whilst at a first glance, our behavioural findings suggest that predictable information is better remembered, an interpretation of findings at the ‘B’ image suggests that the relationship between stimulus predictability and memory encoding may be more nuanced and stimulus dependent.

### 5.2. The relationship between N400 amplitude and memory processing

Whilst our behavioural results highlight the importance of precision and attentional mechanisms in predictive processing, findings relating the N400 to memory performance shed light on the neurophysiological activity that contributes to memory encoding. Notably, we observed a positive relationship between N400 amplitude and memory performance for individuals with a low IAF, whilst this effect was absent for those with a high IAF. Given the well-established sensitivity of the N400 to stimulus expectancy (e.g., DeLong et al., 2005; Wlotko & Federmeier, 2012; for a review, see Kutas & Federmeier, 2011), and it’s links to prediction error (e.g., Bornkessel-Schlesewsky & Schlesewsky, 2019; Fitz & Chang, 2019; Hodapp & Rabovsky, 2021; Rabovsky and McRae, 2014), this may suggest that the extent to which predictability drives memory encoding is subject to individual differences. Prediction error purportedly prompts memory updating via memory reconsolidation mechanisms (Exton-McGuinness et al., 2015; Fernández et al., 2016; Sinclair et al., 2021; Sinclair & Barense, 2019). Reconsolidation is the process by which an existing memory is reactivated, becoming unstable and prone to change, whether that be interference or memory strengthening (Sinclair & Barense, 2019). When a stimulus is encountered, a memory trace for a previously learnt association is reactivated (Kveraga et al., 2007; Sinclair & Barense, 2019). Prediction errors then weaken the associations between the originally learnt items, prompting reconsolidation and enabling the new information to be updated into the existing memory representation (Exton-McGuinness et al., 2015; Sinclair & Barense, 2019). This effect appears contingent on the magnitude of the prediction error, with minimal prediction errors seeming insufficient to prompt reconsolidation and updating of the new, error-inducing information into memory (Gura et al., 2020). In the present study, larger N400 amplitudes may have prompted memory reconsolidation for low IAF individuals, where the unpredictable information was updated into the existing memory trace. However, it is important to explore in further detail why this relationship may differ according to between-subject neural variability.

### 5.3. The effect of N400 amplitude on memory is modulated by individual alpha frequency (IAF)

The present study is the first to our knowledge to demonstrate a role for IAF in modulating the effect of N400 amplitude on memory processing. If the N400 is indeed related to prediction error, this result may imply that the extent to which prediction error promotes learning is subject to between-participant variation. A high IAF has been linked to better general cognitive ability (Grandy, Werkle-Bergner, Chicherio, Lövdén et al., 2013) and to faster information processing speeds, reflected by shorter reaction times (Klimesch et al., 1996; Surwillo, 1961, 1963) and more accurate responses to near-threshold stimuli (Samaha & Postle, 2015). In an emotional memory study by Cross and colleagues’ (2020), high IAF individuals exhibited stronger memory for both neutral and positive items, in contrast to low IAF individuals. Here, negative information was argued to be of greatest importance, and from this it was suggested that high IAF individuals may possess a heightened ability to encode less relevant information into memory (Cross et al., 2020). With this in mind, the weak relationship between the N400 and memory performance in the present study for those with a high IAF may imply that memory encoding can occur independently of the level of prediction error. Whilst it has been previously suggested that high IAF individuals are more likely to update their predictive models (Kurthen et al., 2020), it may also be the case that they adopt a more widespread memory encoding strategy. This potentially explains the positive associations between IAF and memory performance described in previous literature (e.g., Klimesch, 1997; Klimesch et al., 1990, 1993) and implies that the extent to which prediction errors (presumably captured via the N400) drive model updating depends on intrinsic neural differences.

However, it is important to note that the effect of IAF and N400 amplitude on memory performance occurred regardless of whether images belonged to predictable or unpredictable sequences. Although this renders it difficult to confidently conclude that the N400 reflects prediction error in the present study, it does not undermine the likelihood that participants were predicting. The current design, capitalising on statistical regularities in a manner similar to Sherman and Turk-Browne (2020), included consistent categorical relationships that should give rise to predictions (Baker et al., 2014; Bar, 2007; Dale et al., 2012; Mumford, 1992). However, the uniqueness of the items, coupled with the enhanced memory performance and greater N400 amplitudes surrounding the ‘B’ images (as described above), may suggest that our predictability manipulation was less sensitive to subtle differences in item-specific expectancy. To further elucidate the relationship between the N400 and prediction error in future research, a more nuanced manipulation could be introduced. For example, participants could learn contingencies that are later violated (as in Ortiz-Tudela et al., 2023), promoting comparisons between the amplitude of the N400 component and the degree of violation. Additionally, we did not explicitly instruct participants to predict, as doing so would have shifted the focus to explicit forms of prediction, which was not our aim. However, more objective measures of prediction, such as Dale and colleagues’ (2012) analysis of anticipatory computer mouse trajectories, could complement N400 findings by providing converging evidence of predictive processing in future investigations.

Moreover, while the N400 is strongly related to stimulus predictability, it has also been linked to the integration of information with existing meaning representations stored in long term memory (Brown & Hagoort; 1993; Meyer et al., 2007). For low IAF individuals in the current study, difficulty integrating an image with a prior semantic representation may have promoted the encoding of that information as a new memory trace. In contrast, for those with high IAFs, information uptake may have occurred independently of the ease of integration with the prior context. Whilst this study cannot ultimately speak directly to whether integration or prediction underlies this trend, it provides important preliminary evidence demonstrating a relationship between event-related activity, memory updating, and individual neural variability. This could explain inter-subject differences in behavioural memory performance, by suggesting that such variability might stem from disparities in IAF. Interestingly, while the effect of the N400 on memory was modulated by individual variability, the exploratory, late positive component appears to drive learning regardless of IAF.

### 5.4. The significance of late ERP activity for memory encoding

Whilst the observed N400 effects could suggest that predictability influences memory formation for some individuals, differences in the later component (potentially resembling an LPC) may be explained by research on the P300 ERP. In the current research, at the position of the ‘A’ image, the amplitude of this later positivity was greater for unpredictable versus predictable images. In contrast, this effect reversed at the ‘D’ image position. Traditionally, increases in the LPC have been linked to episodic recollection (Curran, 2000; Curran & Cleary, 2003). However, as each image in the learning phase was unique, items seen during this phase could not be explicitly ‘recognised,’ rendering it difficult to explain the observed differences in this component from a purely recollection-based account. However, it is possible that effects here reflect recognition at the category level, as compared to specific episodic recollection, such that LPC amplitudes differed according to recollection of the various image categories.

As an alternative explanation, what we originally deemed a potential LPC might correspond to either a late P300 or a P600. Finnigan and colleagues (2002) discuss how such late positivities are often labelled as an LPC, a P300, or a P600, and for evidence supporting the P600 as a P3 see (Sassenhagen et al., 2014; Sassenhagen & Fiebach, 2019). The P300 component is modulated by stimulus probability and relevance (Berlad & Pratt, 1995; Polich & Margala, 1997), and is commonly observed in oddball paradigms, in response to deviant target stimuli that are presented amongst standard stimuli (for a review see Picton, 1992). For this reason, classical conceptions of the component link it to context updating (i.e., a need to revise one’s schema of the current context following a contextual change; Donchin, 1981). This notion of ‘context updating’ may bear similarities to predictive model updating, potentially explaining the differences in amplitude according to image position and condition observed here. However, it is important to note that the effect of this late component on memory occurred independently of image condition and position, suggesting that these position- and condition-specific amplitude changes do not directly translate to behavioural effects. This could arise because subtle amplitude differences are less pronounced when related to aggregate memory performance measures (such as d’). Nevertheless, this finding, coupled with the P300’s link to context updating (Donchin, 1981), could, at a first glance, support the notion that prediction errors promote memory encoding.

Building upon the link between the late positive ERP observed here and memory formation, recent research demonstrates a correlation between the amplitude of the P600 to semantic violations, and subsequent behavioural recognition memory performance (Contier et al., 2024). Contier and colleagues (2024) interpret these findings in light of the proposal that the N400 reflects implicit prediction error, which in turn drives implicit learning (Hodapp & Rabovsky, 2021; Rabovsky & McRae, 2014). In contrast, the authors link the P600 to explicit memory and highlight the component’s association with sustained attention and conscious awareness (Batterink & Neville, 2013; Contier et al., 2022, 2024). This interpretation also aligns with the P300’s emergence primarily in response to task-relevant or attended-to stimuli (for a review see Picton, 1992), suggesting that such positivities are connected to explicit processes. Furthermore, Bornkessel-Schlesewsky and Schlesewsky (2019) propose that positive language-related ERPs (i.e., the P600 and P300) are response-locked (as opposed to stimulus-locked), and as such reflect behavioural outcomes of predictive coding rather than the process of predictive coding itself (Bornkessel-Schlesewsky & Schlesewsky, 2019).

If the late positive component observed here belongs to the P300/P600 family, the effect of this later component on memory performance could stem from more explicit processes that can additionally drive memory formation. It may also explain why the ERP exhibits a slightly different pattern to the N400 – while the N400 is more reflective of implicit prediction error (Rabovsky et al., 2018; Rabovsky & McRae, 2014), the P600 may relate to explicit processes. Therefore, the possible distinction between the N400 and later, P600-like components suggests that, in the present study, the N400 may better capture the prediction errors arising from implicit predictions (which is what we aimed to investigate), as opposed to the late positivity. Interestingly, the N400 effect on explicit memory recognition (for some individuals) observed here could hint at an overlap between implicit and explicit processes, which would be an intriguing topic for future ERP research.

### 5.5. Conclusion

The present study investigated the effect of stimulus predictability on memory, with behavioural results providing evidence to suggest that predictable information is encoded into memory more readily than unpredictable information. At a first glance, this opposes the notion that predictions, due to their reliance on prior memory representations, inhibit the encoding of new information (Sherman et al., 2022; Sherman & Turk-Browne, 2020). However, the positive correlation between the magnitude of the N400 and subsequent memory for low IAF individuals may be supportive of a prediction error-based learning mechanism, highlighting the complexity of the relationship between prediction and memory. The enhanced memory for predictable ‘B’ images, and the increased N400 amplitudes following items in this position may also imply that memory encoding during prediction is sensitive to subtle differences that surpass basic categorical boundaries of predictability. Importantly, further research would be ideal to determine the extent to which the N400 and the late positivity relate to implicit/explicit prediction and learning. Nevertheless, this study highlights the functional relevance of the two correlates for memory processing, whilst emphasising the importance of individual neural factors for learning.

### Declaration of Generative AI and AI-assisted technologies in the writing process

During the preparation of this work the authors used Chat-GPT sparingly to assist with the refinement of sentences supplied by the authors. After using this tool/service, the authors reviewed and edited the content as needed and take full responsibility for the content of the publication.

## Funding

This research did not receive any specific grant from funding agencies in the public, commercial, or not-for-profit sectors.

## Competing interests

The authors have no competing interests to declare.

## 6. Appendix A

### Statistical model outputs

#### 6.1. Model 1: dprime ∼ condition * cog_scale + condition * slope_scale + (1|subjnr), data= behav_indiv

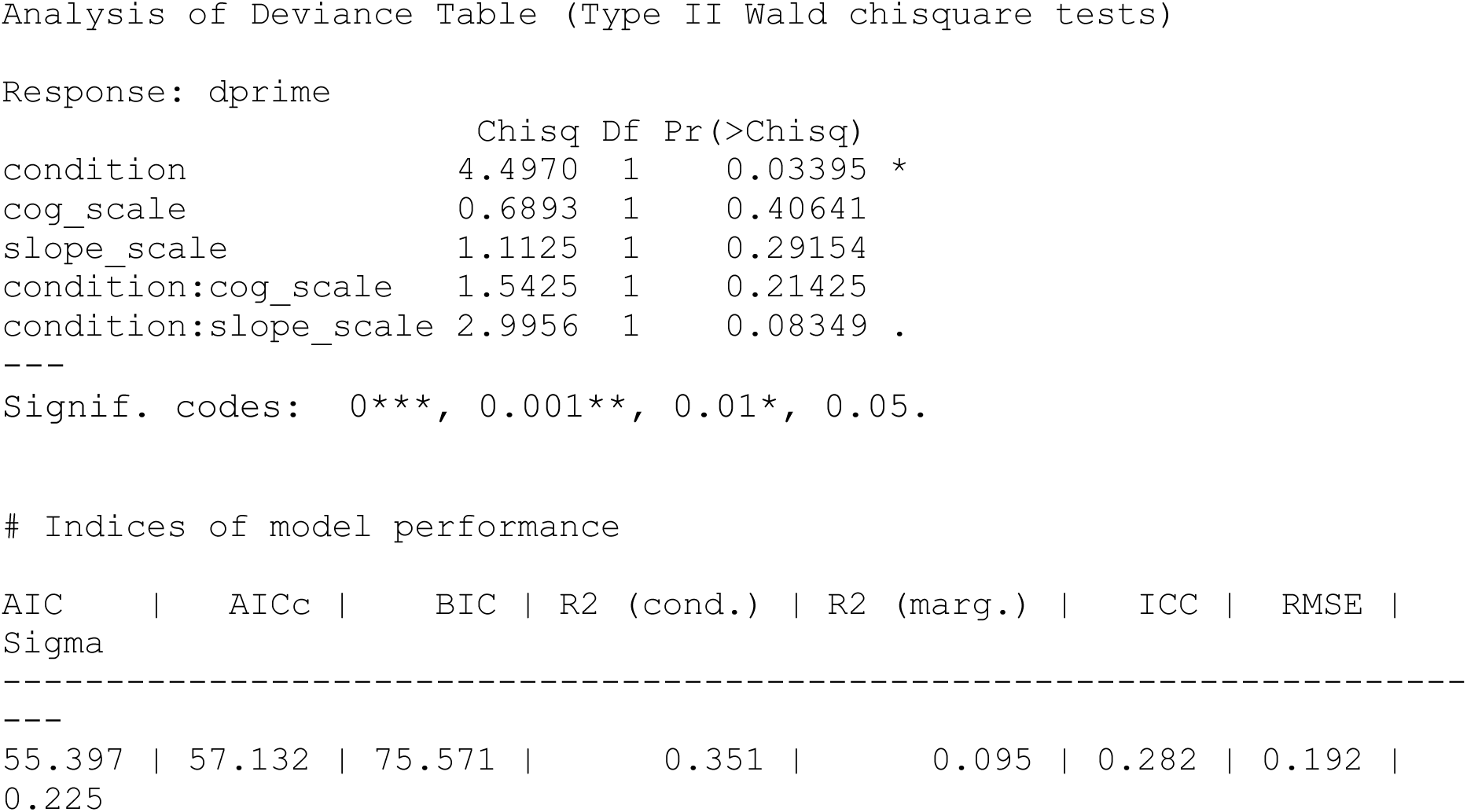

#### 6.2. Model 2: dprime ∼ imagetype * cog_scale + imagetype * slope_scale + (1|subjnr), data= behav_indiv

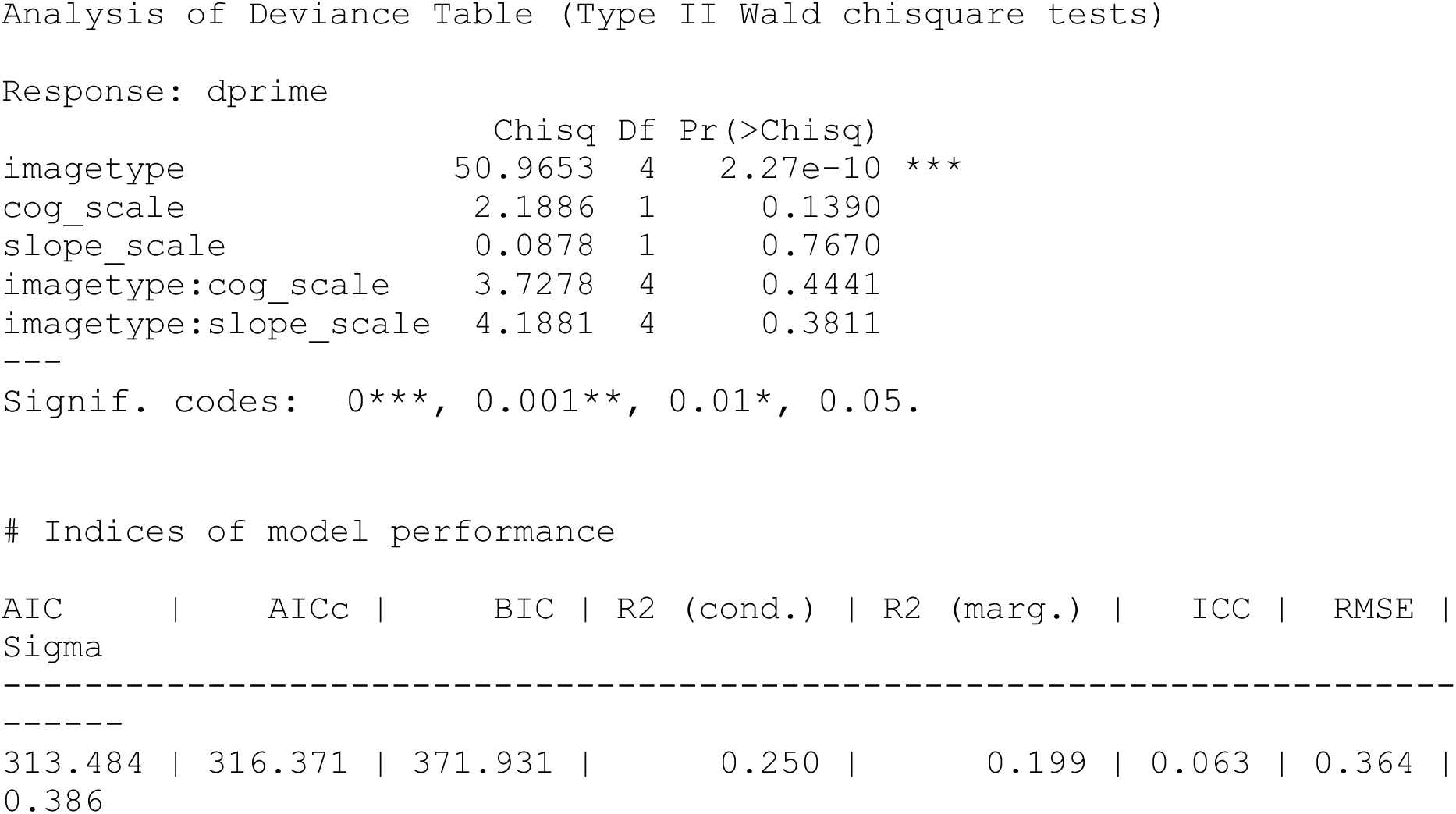

#### 6.3. Model 3: N400amp_mean ∼ sag. * lat. + prestim_amp_scale + (1|subjnr) + (1|item_no), data= N4_small

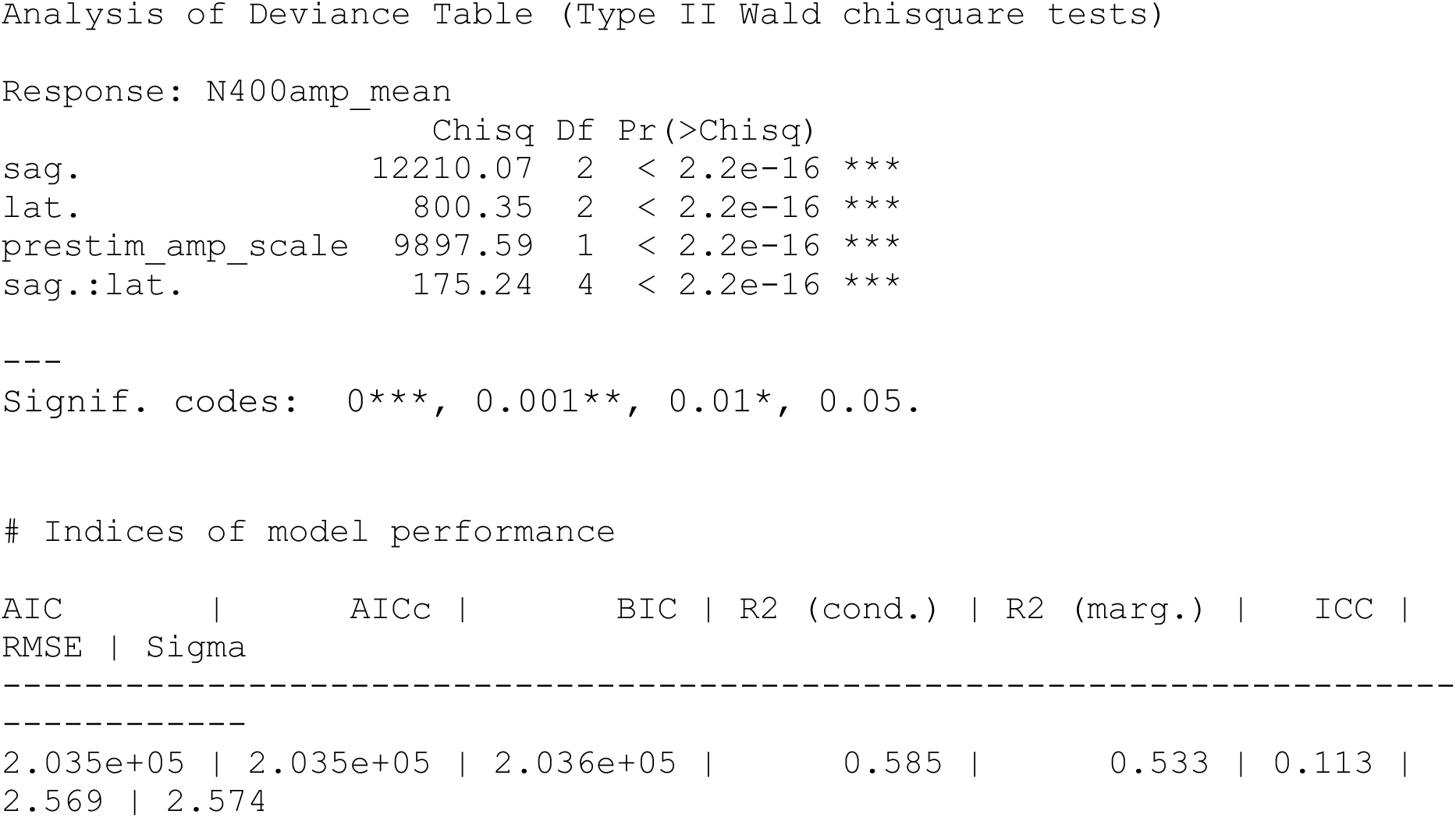

#### 6.4. Model 4: N400amp_mean ∼ condition * position + prestim_amp_scale + (1|subjnr) + (1|item_no), data= N4_region_sum

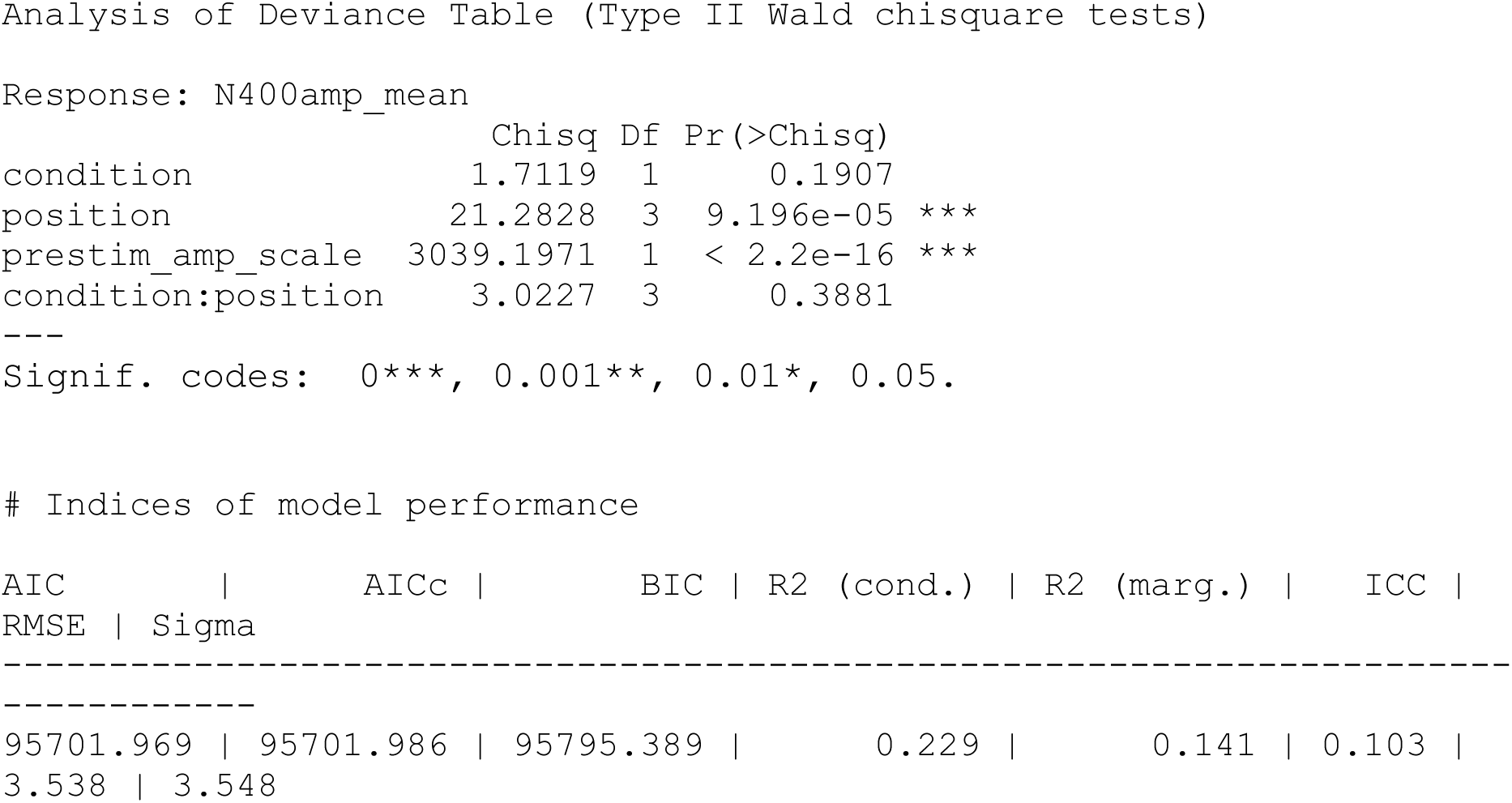

#### 6.5. Model 5: dprime ∼ condition * position * N400amp_scale * cog_scale + prestim_amp_scale + (1+condition|subjnr), data= N4_region_small

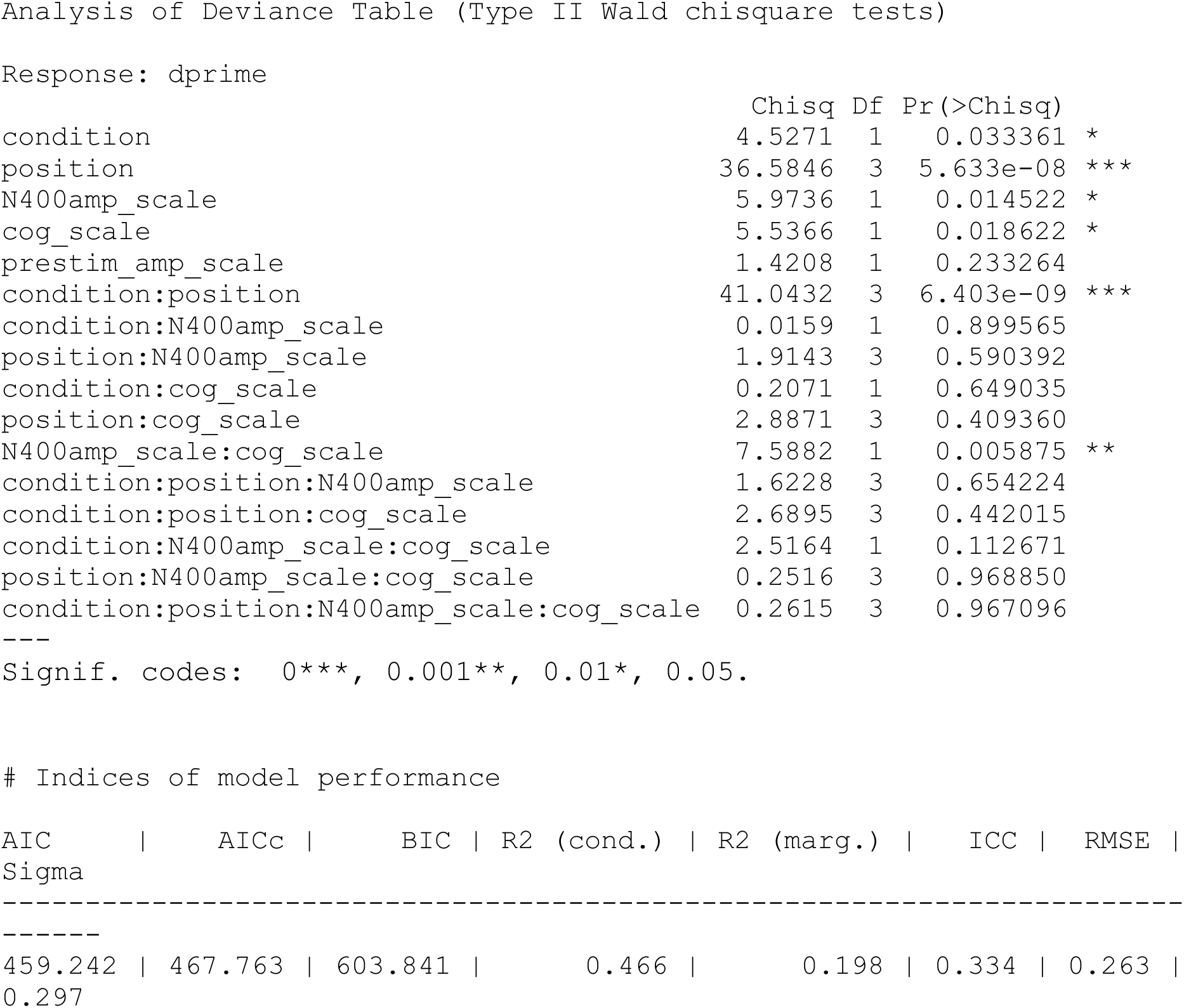

#### 6.6. Model 5.2: dprime∼ condition * position * N400amp_scale * slope_scale + prestim_amp_scale + (1+condition|subjnr), data= N4_region_small

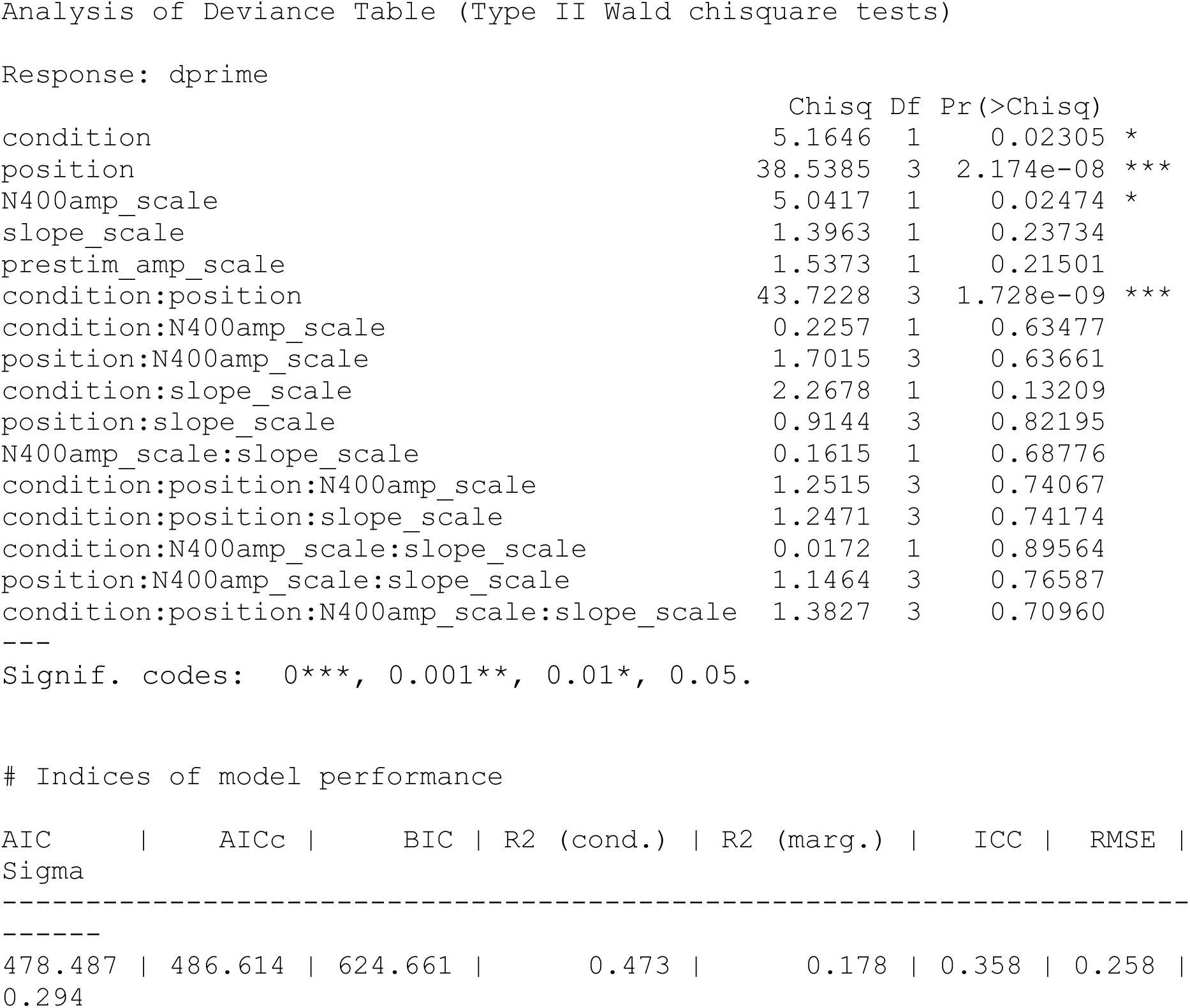

#### 6.7. Model 6: LPCamp_mean ∼ sag. * lat. + prestim_amp_scale + (1|subjnr) + (1|item_no), data= LPC_small

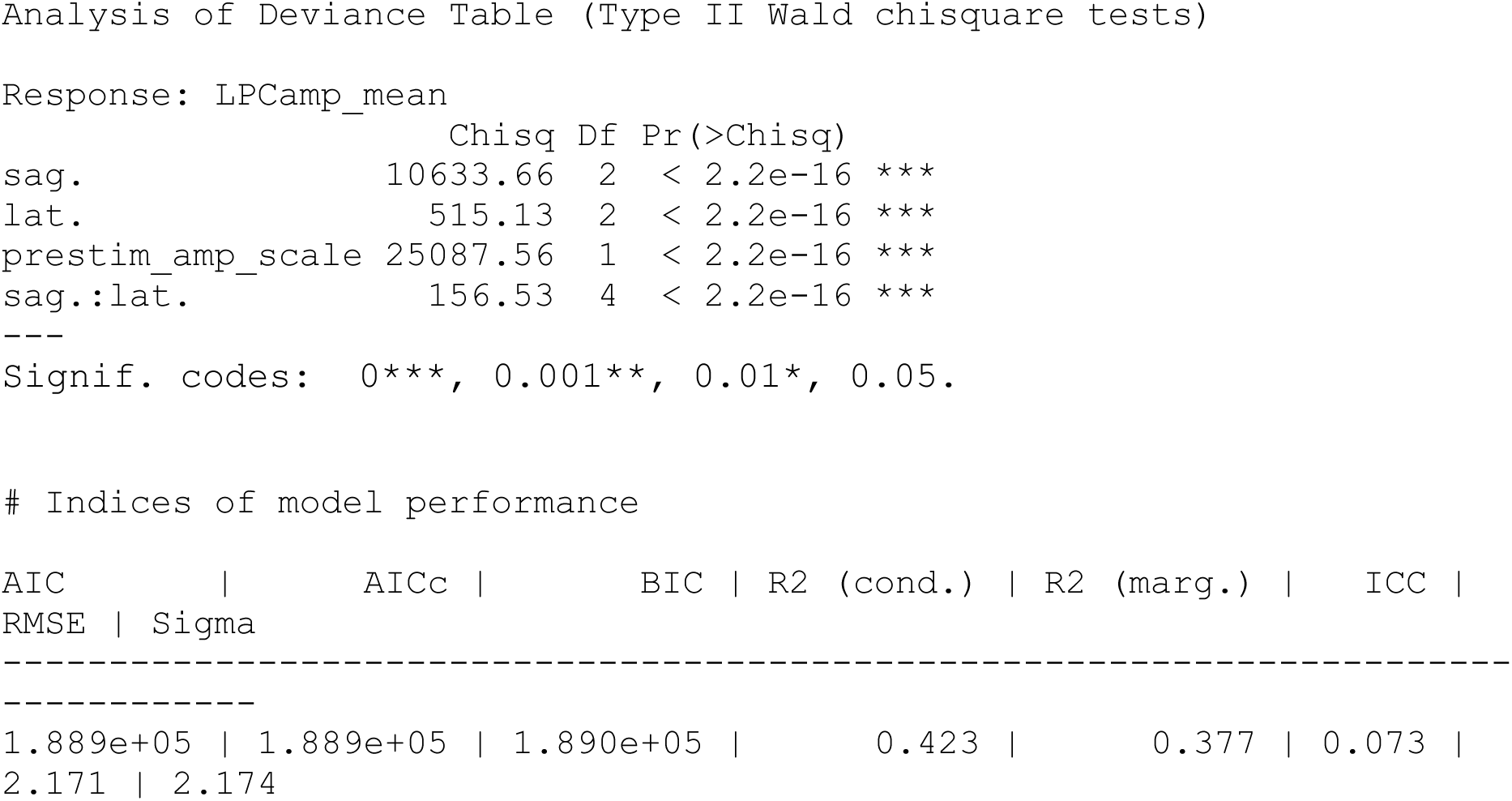

#### 6.8. Model 7: LPCamp_mean ∼ condition * position + prestim_amp_scale + (1+condition|subjnr) + (1|item_no), data= LPC_region_sum

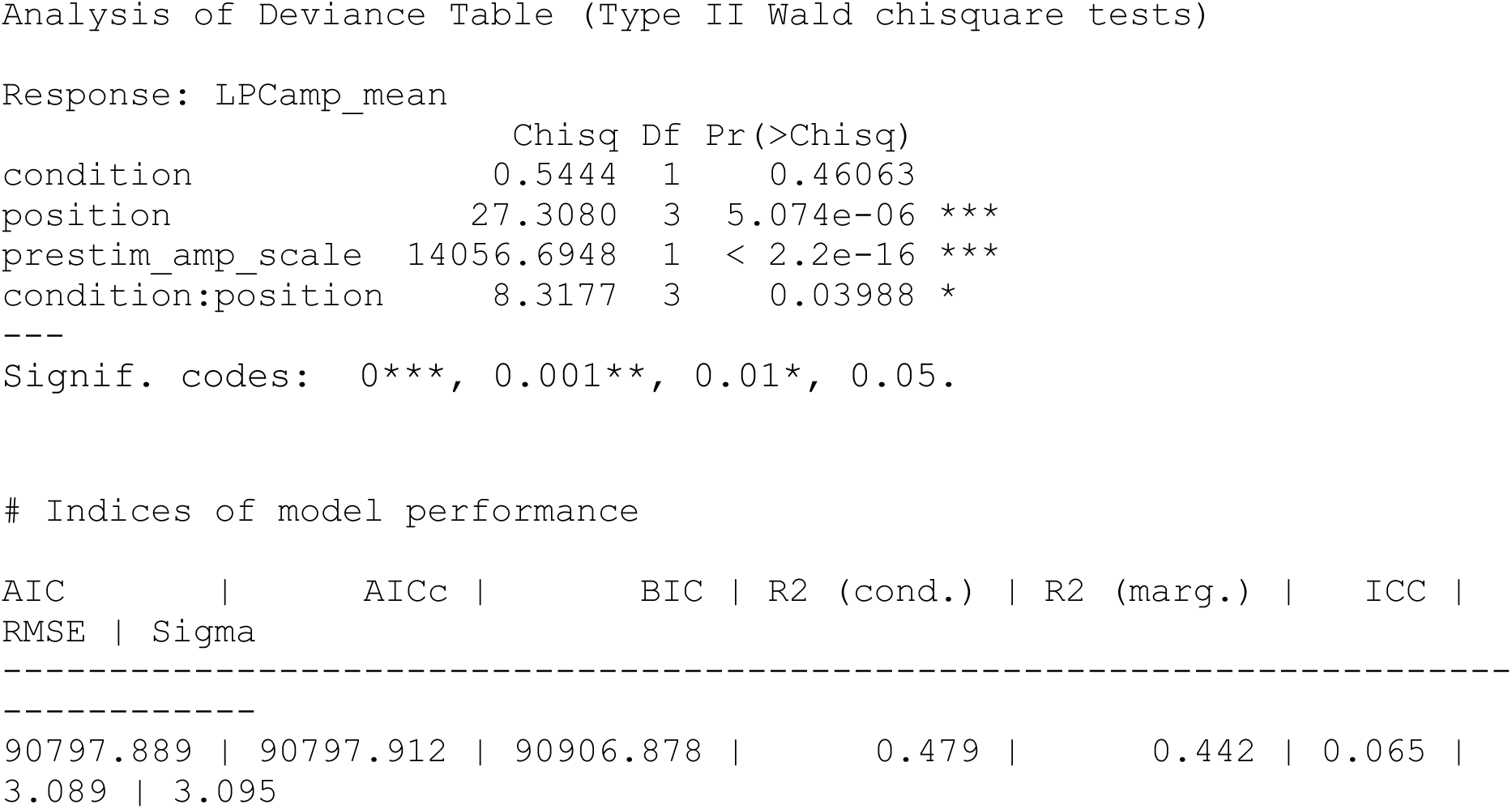

#### 6.9. Model 8: dprime ∼ condition * position * LPCamp_scale + prestim_amp_scale + (1+condition|subjnr), data= LPC_region_small

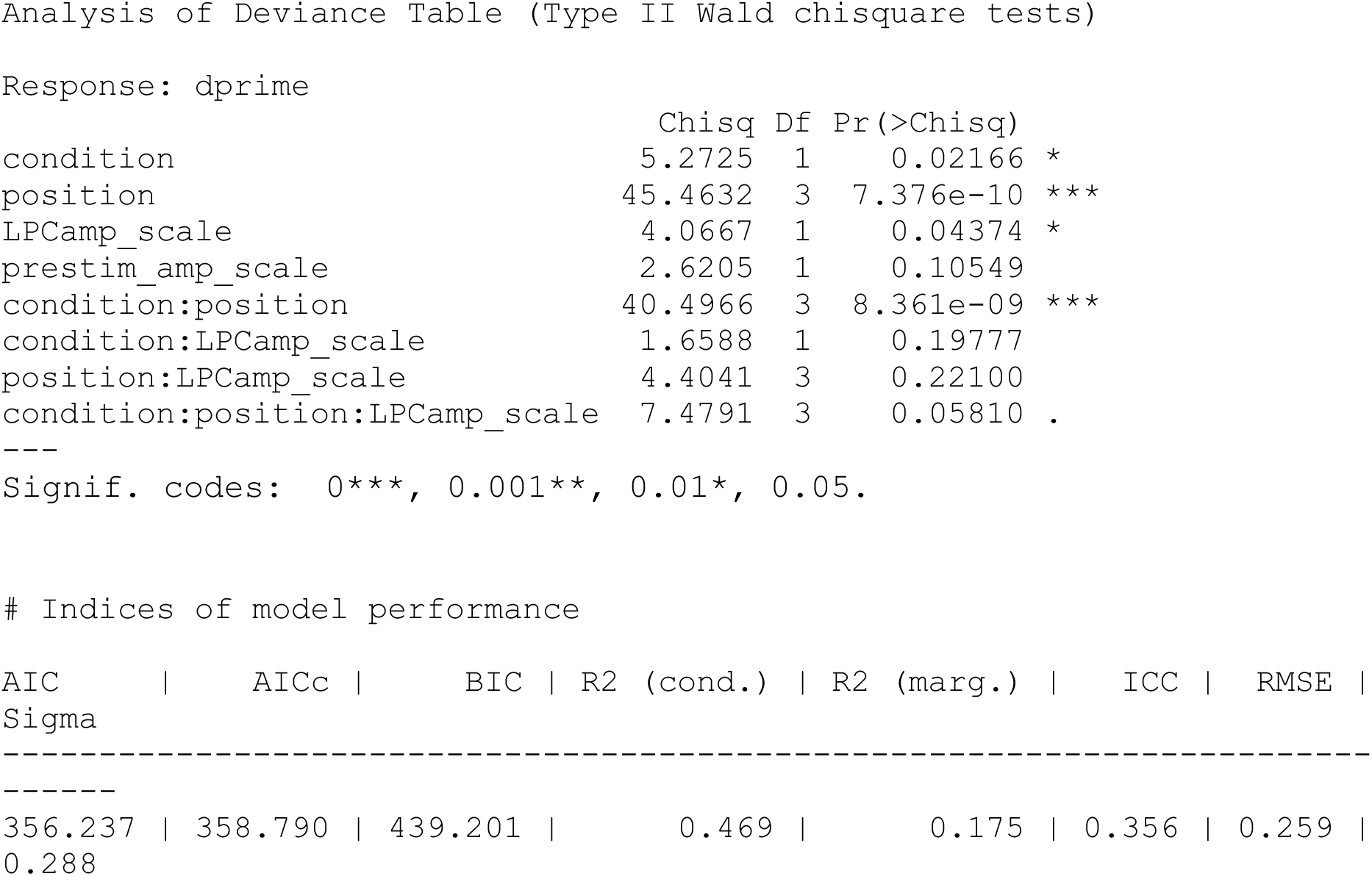

## Notes

https://osf.io/qvjer/

### Competing Interest Statement

The authors have declared no competing interest.

### Summary of Updates

Changes to ERP grand average plots and to Figure 1, an additional analysis and discussion of a later ERP component has been re-incorporated, changes to wording throughout, more clarification and suggestions for future research have been added to discussion

